# Endothelial integration of mechanosensory signals by the spectrin cytoskeleton

**DOI:** 10.1101/2021.08.29.458104

**Authors:** Sivakami Mylvaganam, Bushra Yusuf, Ren Li, Chien-Yi Lu, Lisa A. Robinson, Spencer A. Freeman, Sergio Grinstein

## Abstract

Physiological blood flow induces the secretion of vasoactive compounds, notably NO, and promotes endothelial cell elongation and reorientation parallel to the direction of applied shear. How shear is sensed and relayed to intracellular effectors is incompletely understood. We demonstrate that an apical spectrin network is essential to convey the force imposed by shear to endothelial mechanosensors. By anchoring CD44, spectrin modulates the cell surface density of hyaluronan, which senses and translates shear into changes in plasma membrane tension. Spectrins also regulate the stability of apical caveolae, where the mechanosensitive Piezo1 channels are thought to reside. Accordingly, shear-induced Piezo1 activation and the associated calcium influx were absent in spectrin-deficient cells. As a result, cell realignment and flow-induced eNOS stimulation were similarly dependent on spectrin. We concluded that the apical spectrin network is not only required for shear sensing, but transmits and distributes the resulting tensile forces to mechanosensors that elicit protective and vasoactive responses.

## Introduction

Precise control of the blood flow is required to efficiently perfuse all tissues (Copp, 1995; Levick, 1991). This control relies in large part on endothelial cells, which line the vascular lumen and sense the shear exerted by flowing blood (Davies, 1995; Hahn and Schwartz, 2009). In response to shear, endothelial cells secrete vasoactive compounds including nitric oxide (NO) that act on smooth muscle cells, triggering acute changes in vascular tone and/or initiating chronic vascular remodelling to modulate vessel diameter (Davies, 1995; Hahn and Schwartz, 2009). NO also inhibits pro-coagulant factors in the blood from interacting with the vessel wall, thereby preserving blood fluidity. Importantly, in addition to inducing the secretion of vasoactive compounds, physiological laminar blood flow also promotes endothelial cell elongation and reorientation parallel to the direction of applied shear (Dewey et al., 1981). This cellular alignment confers protection against hemodynamic stress and is associated with anti-inflammatory gene expression (Barbee et al., 1994; Barbee et al., 1995; Davies et al., 1984; Hahn and Schwartz, 2009). Loss of shear-induced endothelial alignment and NO production is associated with increased susceptibility to the formation of atherosclerotic lesions (Davies, 2009; Gimbrone et al., 2000) and with defects in vasculogenesis (Baeyens et al., 2016; Homan et al., 2019). These responses are therefore critical to vascular health.

Several mechanisms of endothelial shear sensing and mechanotransduction have been proposed. One widely accepted model implicates a junctional mechanosensory complex involving the adhesion receptor PECAM-1 (CD31), the junctional protein VE-cadherin and the signaling receptor vascular endothelial growth-factor receptor 2 (VEGFR2). According to this model, shear exerts strain across intercellular junctions, activating PECAM-1 which in turn recruits and activates Src-family kinases (Newman, 1997). Evidence suggests that close apposition of VE-cadherin, VEGFR2, and PECAM-1 facilitates the transactivation of VEGFR2 by PECAM-1-associated Src in response to shear (Tzima et al., 2005). Phosphorylated VEGFR2 is then expected to recruit and activate key downstream signaling molecules including PI3K and Akt (Conway et al., 2013; Tzima et al., 2005). However, PECAM-1-independent, shear-induced activation of several signaling pathways has been observed in the endothelium (Jin et al., 2003). Moreover, PECAM-1-deficient mice develop normally and without any notable defects in cardiovascular morphogenesis (Duncan et al., 1999). These observations are inconsistent with an indispensable role for PECAM-1 in regulating endothelial responses to shear (Davies, 2009).

It is likely that other, junction-independent processes play a critical role in endothelial mechanotransduction. In this regard, it is noteworthy that the glycocalyx has been implicated in shear-induced vascular responses. Of the glycocalyx components, hyaluronic acid (HA) seems particularly important. HA is a non-sulfated glycosaminoglycan synthesized at the plasma membrane by HA synthases (HAS1, 2, and 3), integral membrane enzymes that catalyze the transglycosylation of UDP-glucoronate and UDP-N-acetylglucosamine while simultaneously extruding the growing linear polymer into the vascular lumen. The elongating HA polymer stays associated with the cell via the synthases until such time that the polymerization arrests, when it is released (Necas et al., 2008). HA can also be anchored to the cell surface by binding to its receptors, primarily the glycoprotein CD44 (Aruffo et al.1990). Importantly, the enzymatic degradation of vascular HA disrupts normal endothelial responses to shear *in vitro* and *in vivo* (Mochizuki et al., 2003; Pahakis et al., 2007; Tarbell and Ebong, 2008), and inhibiting HA production by genetic deletion of HAS2 causes severe abnormalities in cardiac and vascular morphogenesis, and is embryonically lethal (Camenisch et al., 2000). In addition to the glycocalyx, mechanosensitive ion channels including Piezo1 are also implicated in normal endothelial responsiveness to shear, mediating critical Ca^2+^ signals that elicit downstream reactions (Li et al., 2014). It thus appears that multiple, seemingly independent molecules cooperate to generate the endothelial responses to flow. However, a coherent mechanism describing how these are coordinated is lacking.

We recently identified the presence of a polarized spectrin network restricted to the apical (luminal) aspect of endothelial cells, the surface that most directly experiences fluid shear (Mylvaganam et al., 2020). Spectrin networks are comprised of flexible *α*- and *β*-subunit heterodimers that stabilize short actin filaments at membranes (Bennett and Baines, 2001). Together with its associated proteins, spectrin has been implicated in regulating the dynamics of glycoproteins and surface ion channels in multiple cells (Sheetz et al., 1980; Xu et al., 2013).

Importantly, in the endothelium, the spectrin cytoskeleton controls the apical (luminal) retention and stabilization of CD44, the primary receptor for HA (Mylvaganam et al., 2020). It thus seems plausible that spectrin may play a role in regulating endothelial responses to shear, at least in part through its associations with CD44/HA.

In an effort to resolve the uncertainties regarding the relative contribution of the various endothelial mechanotransduction processes, we used a combination of imaging and biophysical approaches to systematically analyze the role of the intercellular junctional complex, HA and Piezo1 in shear-sensing. We also examined the possibility that spectrin networks may serve as integrators and/or transducers of the forces imposed by flowing fluid.

## Results

### Cell-cell junctions are not required for shear-induced endothelial alignment

Due to the importance of endothelial mechano-responses in the arterial circulation, we used primarily an immortalized line of human aortic endothelial cells (telo-HAEC, or tHAEC) for our analyses. To investigate the mechanisms underlying their responses to shear, we examined the behavior of cells grown in collagen-coated microfluidic chambers and subjected to a constant shear stress of 15 dynes/cm^2^, mimicking conditions encountered in portions of the arterial circulation (Doucette et al., 1992). Using this system, we evaluated changes in endothelial morphology in response to applied shear by fixing and staining the cells with phalloidin to visualize F-actin (Fig. 1A). Consistent with previous studies, if grown under static conditions, endothelial cells in confluent monolayers had actin stress fibers that were oriented randomly (Fig. 1A). After brief (30 min) exposure of the monolayer to flow, the cells elongated in the direction of the applied stress, which was coincident with the parallel orientation of their actin stress fibers (Fig. 1A). We developed an analysis pathway to quantify the degree of actin stress fiber alignment from these images by first segmenting individual fibers using a Hessian-based multiscale filter to identify tubular structures, or “filaments”. The angle between the longest axis (the maximum Feret diameter) of each segment and the direction of applied flow was then calculated (Fig. 1A). The interquartile range of the distribution of segmented filament orientations was used to quantify the variability between conditions and revealed significantly less variation in the F-actin orientations – which we interpreted as alignment – in cells exposed to shear when compared with those in static culture (Fig. 1A).

**Figure 1.**
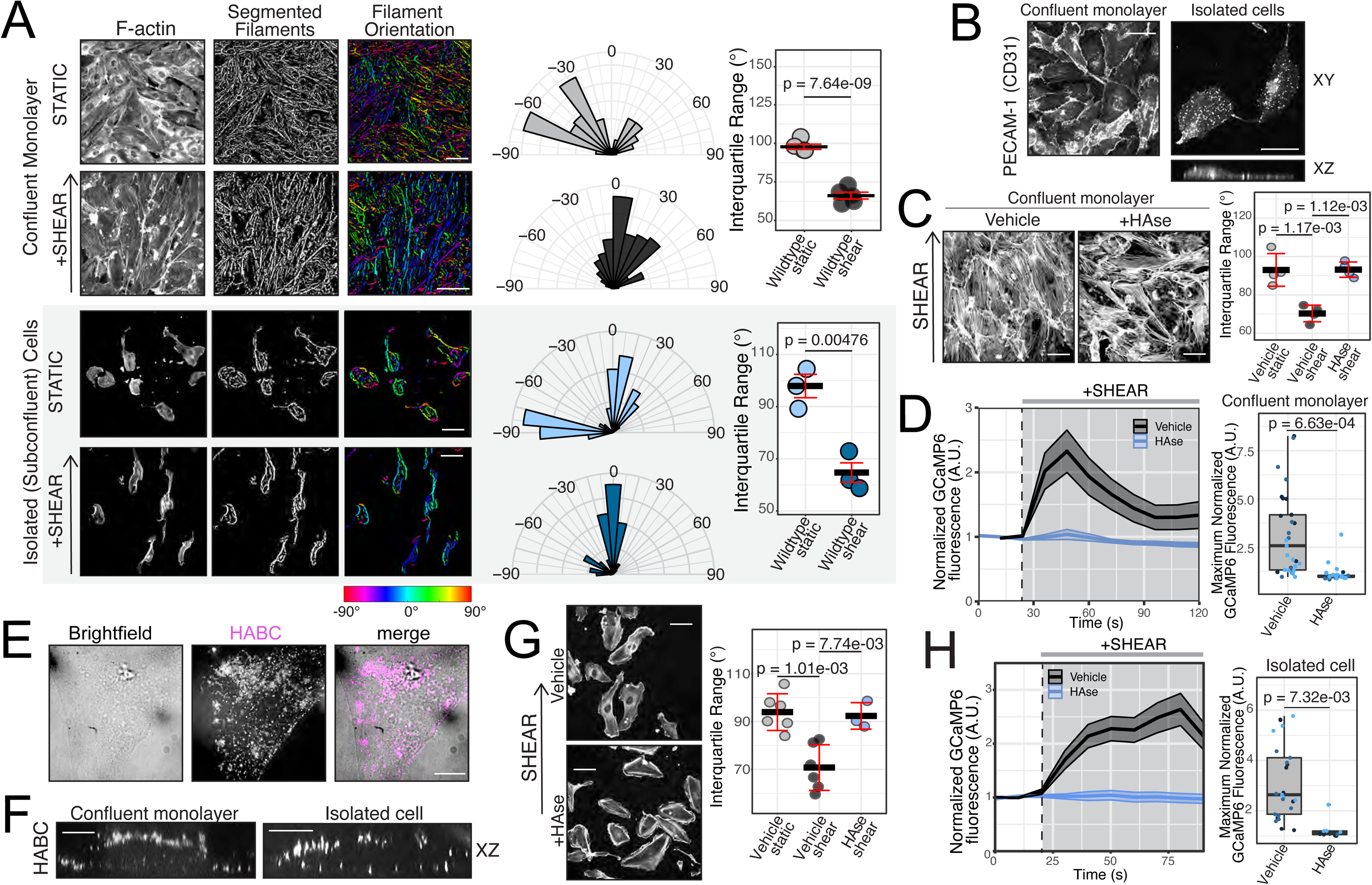
Hyaluronan is an indispensable, junction-independent regulator of endothelial responses to shear stress. **A)** Confluent endothelial monolayers (*top two rows*) or sparsely distributed single endothelial cells (*bottom two rows*) were grown in collagen-coated microfluidic chambers under static conditions or subjected to a constant shear stress of 15 dynes/cm^2^ for 30 min, as indicated prior to fixation and phalloidin staining. *First (leftmost) column*: visualization of F-actin. S*econd column*: images were skeletonized and actin filaments were segmented. *Third column:* color labelling according to orientation of the maximum Feret diameter of the segmented filament relative to the shear axis (0°). *Fourth column:* representative polar histograms of F-actin orientations analyzed as described in Methods from one representative experiment. *Fifth (rightmost) column*: interquartile ranges (the ranges between the 25^th^ to 75^th^ percentile of the data) of the distribution of segmented filament orientations for the indicated conditions. Here and elsewhere, data are means ± SE of 3 independent experiments, each quantifying ≥50 cells. **B)** Distribution of PECAM-1 (CD31) in confluent endothelial monolayers *(left)* and single cells *(right)*, assessed by immunofluorescence. Representative of 3 independent experiments. **C-D)** Confluent monolayers of endothelial cells were treated with vehicle (PBS) or hyaluronidase (HAse) and subjected to constant fluid flow as in 1A. **C)** Representative images of cells fixed and stained with phalloidin (F-actin, *left*) and interquartile ranges of F-actin orientations (*right*). Data are means ± SE of 3 independent experiments, each quantifying ≥50 cells. **D)** Live imaging of endothelial cells expressing the cytosolic [Ca^2+^] indicator GCaMP6 before *(white background)* and after the application of shear *(grey background)*; images were acquired every 12 sec. *Left:* quantification of mean GCaMP6 fluorescence over time, normalized to initial resting value. Data are means ± SE (shaded area, here and elsewhere) of 3 independent experiments, each quantifying ≥8 cells. *Right:* maximum normalized GCaMP6 fluorescence evoked by shear. Here and elsewhere, dots represent maximum values for each cell, color-coded by experiment. **E-F)** Confluent and sparsely distributed subconfluent endothelial cells were incubated with fluorescent HA-binding complex (HABC) and imaged. **G-H)** Sparsely distributed endothelial cells were treated with vehicle (PBS) or hyaluronidase (HAse) and analyzed as in 1C-D. Here and elsewhere, indicated *p*-values are from statistical analyses of experimental means.

Intriguingly, the equivalent treatment of sub-confluent endothelial cultures revealed that isolated cells lacking visible intercellular contacts retained the ability to elongate and align following exposure to shear stress, mimicking the behavior observed for endothelial monolayers (Fig. 1A). This occurred despite the internalization of PECAM-1 from the surface of single cells (Fig. 1B). Sequestration of these proteins into intracellular compartments is expected to preclude their mechanosensory functions. These observations therefore imply that the junctional mechanosensory complex plays a dispensable role in the alignment response of endothelial cells to shear. Other mechanosensors are therefore likely to mediate endothelial alignment in response to shear.

### Junction-independent responses to shear require hyaluronic acid

We considered the contributions of other proposed mechano-sensors to junction-independent responses to shear. We proceeded to examine the role of HA in mediating shear-induced F-actin alignment and Ca^2+^ signaling. Endothelial cells respond to acute changes in their mechanical environment with rapid increases in their cytosolic concentration of free calcium ([Ca^2+^]_cyto_) (Buga et al., 1991; Shen et al., 1992), which are critical in orchestrating cellular alignment and NO production by the endothelium (Buga et al., 1991; Lückhoff et al., 1988). During these signaling events, [Ca^2+^]_cyto_ increases proportionally to the magnitude of the applied force (Hong et al., 2006; Uematsu et al., 1995), thus serving as a clear indicator of shear sensing.

We first confirmed that the enzymatic degradation of surface HA with hyaluronidase (HAase) resulted in the loss of shear-induced F-actin alignment of endothelial cell monolayers (Fig. 1C). To examine the mechanisms governing the elevation of endothelial [Ca^2+^]_cyto_ in response to shear, we monitored changes in the fluorescence of the genetically-encoded cytosolic Ca^2+^ indicator, GCaMP6s in response to various stimuli. Consistent with previous studies (Mochizuki et al., 2003), the application of flow-induced shear stress to otherwise untreated (control) cells was associated with a rapid, pronounced increase in cytosolic [Ca^2+^] (indicated by an increase in GCaMP6s fluorescence) that returned to near-baseline levels within minutes despite continuous exposure to fluid flow (Fig. 1D). Remarkably, pretreating the cells with HAase virtually eliminated the shear-induced [Ca^2+^]_cyto_ transient (Fig. 1D).

To investigate whether HA also regulates shear-sensing in isolated endothelial cells, we first evaluated whether HA was retained at the surface of cells lacking intercellular contacts. We visualized the distribution of HA on the endothelial surface using a biotinylated HA-binding complex (HABC) labelled with fluorophore-conjugated streptavidin (Fig. 1E, 1F). Analysis of live, subconfluent cells revealed extensive regions with dense HABC staining (Fig. 1E). Furthermore, as in confluent cells, HABC bound predominantly to the apical surface of isolated endothelial cells (Fig. 1F). HA was thus localized to the surface of endothelial cells that most directly experiences shear. Indeed, HAase treatment eliminated the shear-induced alignment of sparsely distributed cells to the same extent as it inhibited the alignment of confluent monolayers (Fig. 1C,G). The isolated cells also required HA for the application of shear stress to elicit a [Ca^2+^]_cyto_ elevation (Fig. 1H). We therefore concluded that in addition to its previously described roles in confluent endothelial tissues, HA is able to mediate junction-independent cellular responses to fluid flow.

### CD44, the primary HA receptor, regulates shear-induced alignment *in vitro*

When exerted on the glycocalyx, forces like fluid shear are expected to rapidly distribute throughout this gel-like substance (Weinbaum et al., 2007). Core glycoproteins and glycosaminoglycans that are embedded in this layer are displaced by this force and can then, in theory, deliver information about the direction and magnitude of the applied stress to other membrane-associated proteins (Davies, 1995). Because HA is not directly attached to the plasma membrane, the previous observations implied that receptors connecting the glycopolymer to the endothelium may be important in relaying the mechanical signal. We were particularly interested in CD44, the primary receptor for HA in most tissues (Aruffo et al., 1990).

By immunofluorescence, we observed that CD44 accumulated in the apical plasma membrane of sparsely distributed endothelial cells (Fig. 2A), as found earlier in confluent monolayers. It was therefore plausible that it may help establish and maintain a sufficient density of HA on the surface of isolated cells (Fig. 1E, 1F). Accordingly, genetic depletion of CD44 expression by RNA interference (S. Fig. 1) was associated with defective responses to shear (Fig. 2B). This indicated that CD44 was required for normal shear responses *in vitro*. More broadly, it appeared that proteins that bridge glycocalyx components to the endothelial surface play an important role in its mechanosensory behavior.

**Figure 2.**
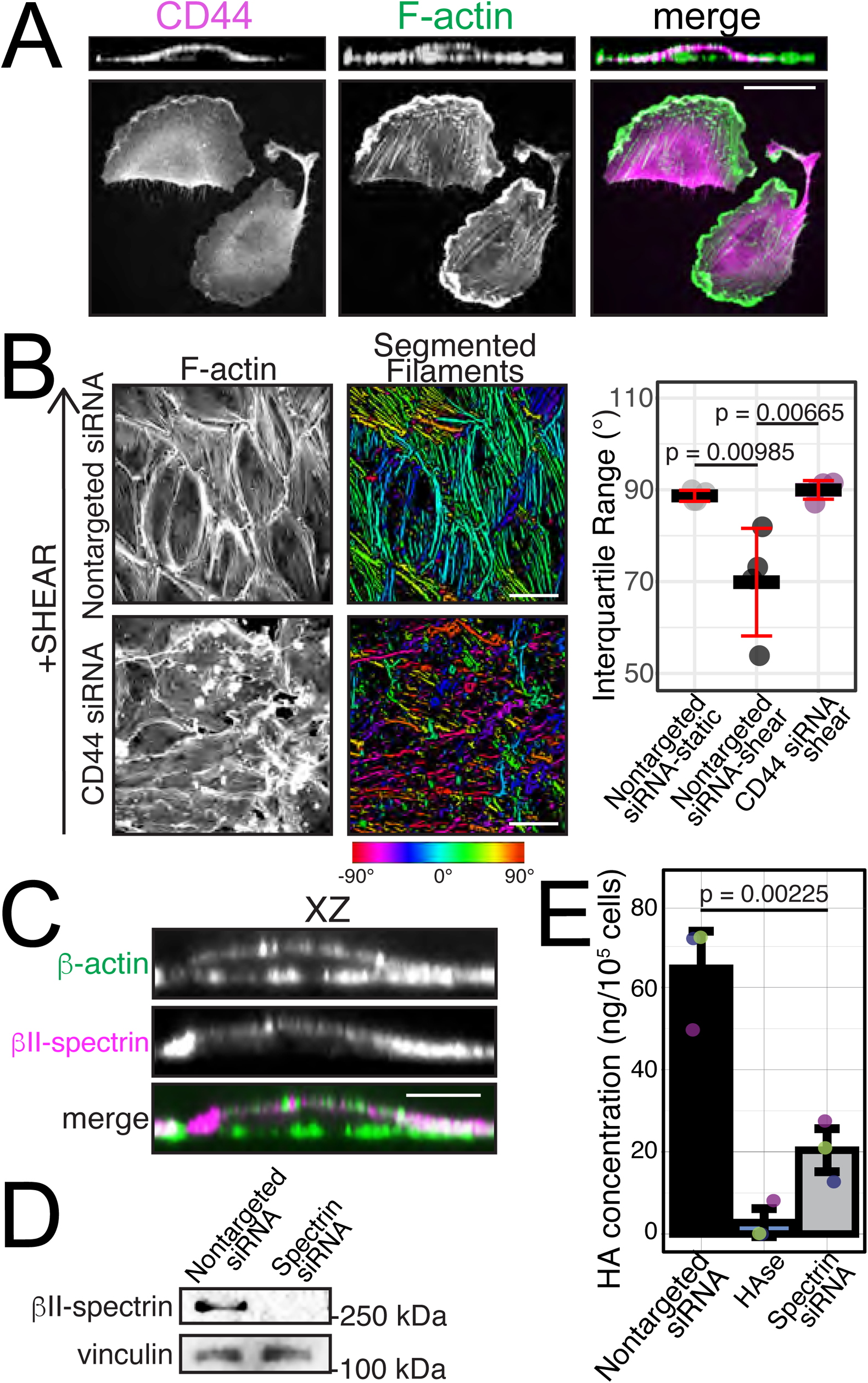
The hyaluronan receptor CD44 mediates shear-induced alignment. **A)** Representative confocal images of sparsely distributed endothelial cells immunostained for endogenous CD44 (*left panel and magenta in merge*) and stained for actin filaments (F-actin) with phalloidin (*middle panel and green in merge*). Representative of 3 independent experiments. **B)** Confluent monolayers of endothelial cells were transfected with non-targeted or CD44 siRNA and subjected to constant fluid flow as in 1A. Representative images of cells fixed and stained with phalloidin (F-actin, *left*), which were then segmented according to filament orientation (*middle). Right panel*: interquartile ranges of F-actin orientations provided. Data are means ± SE of 3 independent experiments, each quantifying ≥50 cells. **C)** Representative orthogonal confocal section of endothelial cell stained for F-actin with phalloidin (*top panel and green in merge*) and immunostained for endogenous *β*II-spectrin (*middle panel and magenta in merge*). **D)** *β*II-spectrin expression in RF24 cells probed by immunoblotting after transfection with non-targeted or *β*II spectrin-targeted siRNAs. Vinculin was probed for normalization. **E)** Concentration of HA on the endothelial surface under the indicated conditions, as measured by ELISA. Individual experiments colored differently. Data are means ± SE of 3 independent experiments.

### Spectrin is required to maintain HA on the luminal endothelium

It remained unclear how transmembrane glycoproteins like CD44 activate downstream signaling cascades. We had previously observed that the apical retention and immobilization of CD44 are regulated by the spectrin cytoskeleton (Mylvaganam et al., 2020), which is predominantly localized to the apical surface in endothelial cells (Fig. 2C). As an extensive cytoskeletal network that spans the apical plasma membrane (Fig. 2C), we considered the possibility that spectrin might facilitate communication between the glycocalyx and other plasmalemmal molecules.

We investigated the potential relationship between HA and spectrin by studying whether HA abundance is normal on the surface of *β*-spectrin-knockdown endothelial cells (Fig. 2D). As noted, α/β spectrin heterodimers are the functional units of the spectrin cytoskeleton and depleting either subunit is therefore an effective strategy for preventing the assembly of these networks. Wild-type endothelial cells grown in culture had appreciable surface HA that was detectable by enzyme-linked immunosorbent assay (ELISA) and was virtually eliminated by HAase (Fig. 2E). Importantly, spectrin-depletion by RNA interference was associated with a pronounced decrease in the surface density of HA (Fig. 2E). These experiments thus provided suggestive evidence that spectrin may regulate endothelial functions associated with HA.

### Spectrin is required for normal endothelial responses to shear

To perform a systematic analysis of the role of spectrin in mediating endothelial responses to shear, we edited the tHAEC cell line to effectively knock out *β*II-spectrin using CRISPR-Cas9 (Fig. 3A). *β*II-spectrin-knockout (KO) endothelial cells did not exhibit any overt morphological defects and the distribution of junctional-complex proteins to intercellular contacts in confluent monolayers was comparable to that observed in wild-type cells (Fig. 3B). Ratios of VE-cadherin fluorescence intensity in the junctional membrane to that in the cytosol were also similar between the KO clones and the parental wild-type cells, indicating that the loss of *β*II-spectrin was not associated with overt defects in junctional integrity (S. Fig. 2A).

**Figure 3:**
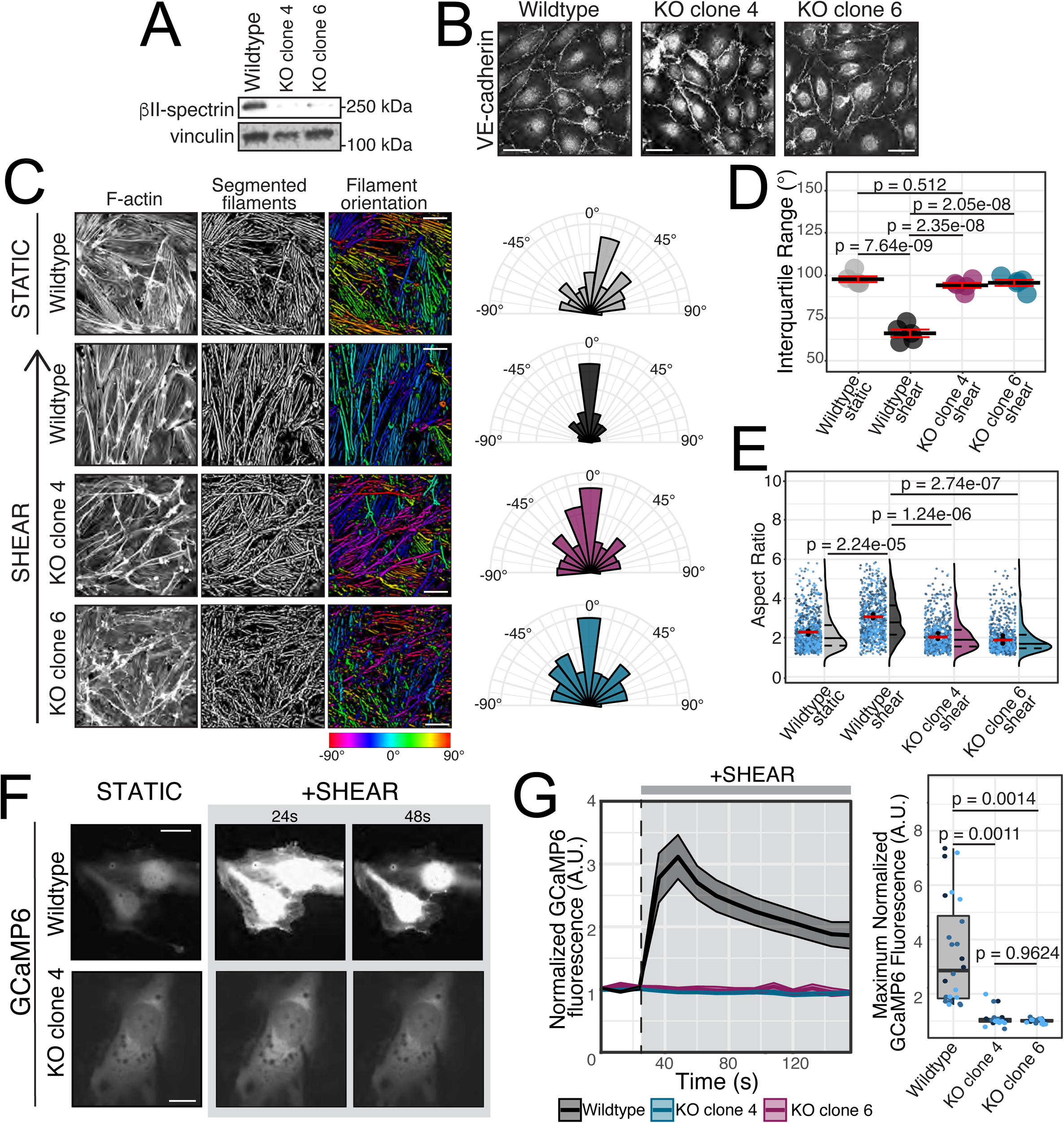
Endothelial cells require spectrin for normal responses to shear stress. **A)** *β*II-spectrin expression in wildtype or spectrin KO tHAEC cell lines generated using CRISPR/Cas9 technology (see methods). Representative of 3 independent experiments. **B)** Distribution of VE-cadherin in indicated endothelial cell lines, assessed by immunofluorescence. Representative of 3 independent experiments. **C)** Endothelial monolayers were grown in collagen-coated microfluidic chambers under static conditions (top) or subjected to a constant shear stress as in **1A**. *First (leftmost) column*: visualization of F-actin. S*econd column*: images were skeletonized and actin filaments were segmented. *Third column:* color labelling according to orientation of the maximum Feret diameter of the segmented filament relative to the shear axis (0°). *Fourth column:* representative polar histograms of F-actin orientations analyzed as described in Methods from one representative experiment. **D)** Interquartile ranges (the ranges between the 25^th^ to 75^th^ percentile of the data) of the distribution of segmented filament orientations for the indicated conditions. Data are means ± SE of 3 independent experiments, each quantifying ≥50 cells. **E)** Confluent endothelial cells grown in static or shear conditions were fixed and immunostained for VE-cadherin to visualize cell borders. Aspect ratios were calculated as the longest axis divided by the shortest axis of each cell. Here and elsewhere, for each condition, on the left: jittered data points (in shades of blue) represent individual measurements and are color-coded according to biological replicate. Means of individual experiments are presented in black. Overall means ± SE are indicated in red. Histograms of all data points indicating the mean (solid line) as well as the 25^th^ and 75^th^ percentiles (dashed line) are shown to the right of the individual data. Data from 3 independent experiments. **F-G)** Live imaging of wildtype (top) and spectrin KO endothelial cells (bottom) expressing the cytosolic [Ca^2+^] indicator GCaMP6 before and after the introduction of shear at 24 sec (grey background). **F)** Representative images acquired before (left) and immediately after application of shear and a further 24 sec later are shown. **G)** *Left:* quantification of mean GCaMP6 fluorescence over time, normalized to initial resting value. Data are means ± SE of 3 independent experiments, each quantifying ≥8 cells. *Right:* maximum normalized GCaMP6 fluorescence evoked by shear.

Remarkably, following exposure to fluid flow for 30 min, stress fiber alignment failed to occur in *β*II-spectrin-KO cells (Fig. 3C, 3D). Instead, the architecture of F-actin networks in these cells more closely resembled that observed in wild-type cells in static culture (Fig. 3C, 3D). Moreover, in addition to alignment, shear is also known to induce the elongation of endothelial cells along the flow axis (Dewey et al., 1981). Changes in cell shape were assessed by immunostaining the junctional protein VE-cadherin in cells subjected to static or shear-stress conditions. Cellular aspect ratio –the ratio of the long axis to the short axis of each cell– was then analyzed from these images. As expected, aspect ratios of wild-type tHAEC monolayers following exposure to shear stress were significantly higher than those of cells maintained in static conditions, indicating a more elongated phenotype (Fig. 3E). Aspect ratios of *β*II-spectrin-KO cells exposed to shear stress were comparatively lower, indicating a failure to elongate (Fig. 3E). Furthermore, no appreciable changes in [Ca^2+^]_cyto_ were detected in *β*II-spectrin-KO cells in response to shear (Fig. 3F, 3G). On the other hand, shear-induced increases in tyrosine phosphorylation of junctional proteins were comparable between wild-type and *β*II-spectrin-KO cells (S. Fig. 2B, 2C). We concluded that spectrin was required for shear-induced Ca^2+^ responses and the associated changes in endothelial morphology, independently of junctional complex proteins.

### Piezo1 activation is regulated by spectrin

The shear-induced elevation of [Ca^2+^]_cyto_ in endothelial cells requires extracellular Ca^2+^ (Schwarz et al., 1992; Shen et al., 1992), implicating channels localized to the plasma membrane that are directly or indirectly mechanosensitive. Multiple shear-sensitive Ca^2+^ channels have been proposed to participate. Shear triggers the release of ATP, which then acts in an autocrine manner to induce Ca^2+^ entry. P2X4 purinoceptors, that are ATP-operated cation channels (Yamamoto et al., 2000), and P2Y receptors, a family of nucleotide-activated G-protein coupled receptors (Wang et al., 2015), have both been implicated in endothelial mechanotransduction. However, we found that ATP-induced Ca^2+^ transients in *β*II-spectrin-KO cells were comparable to those observed in wild-type cells (Fig. 4A). This indicated that the defective responses to shear in spectrin-deficient cells were not attributable to impaired purinergic receptor activity.

**Figure 4:**
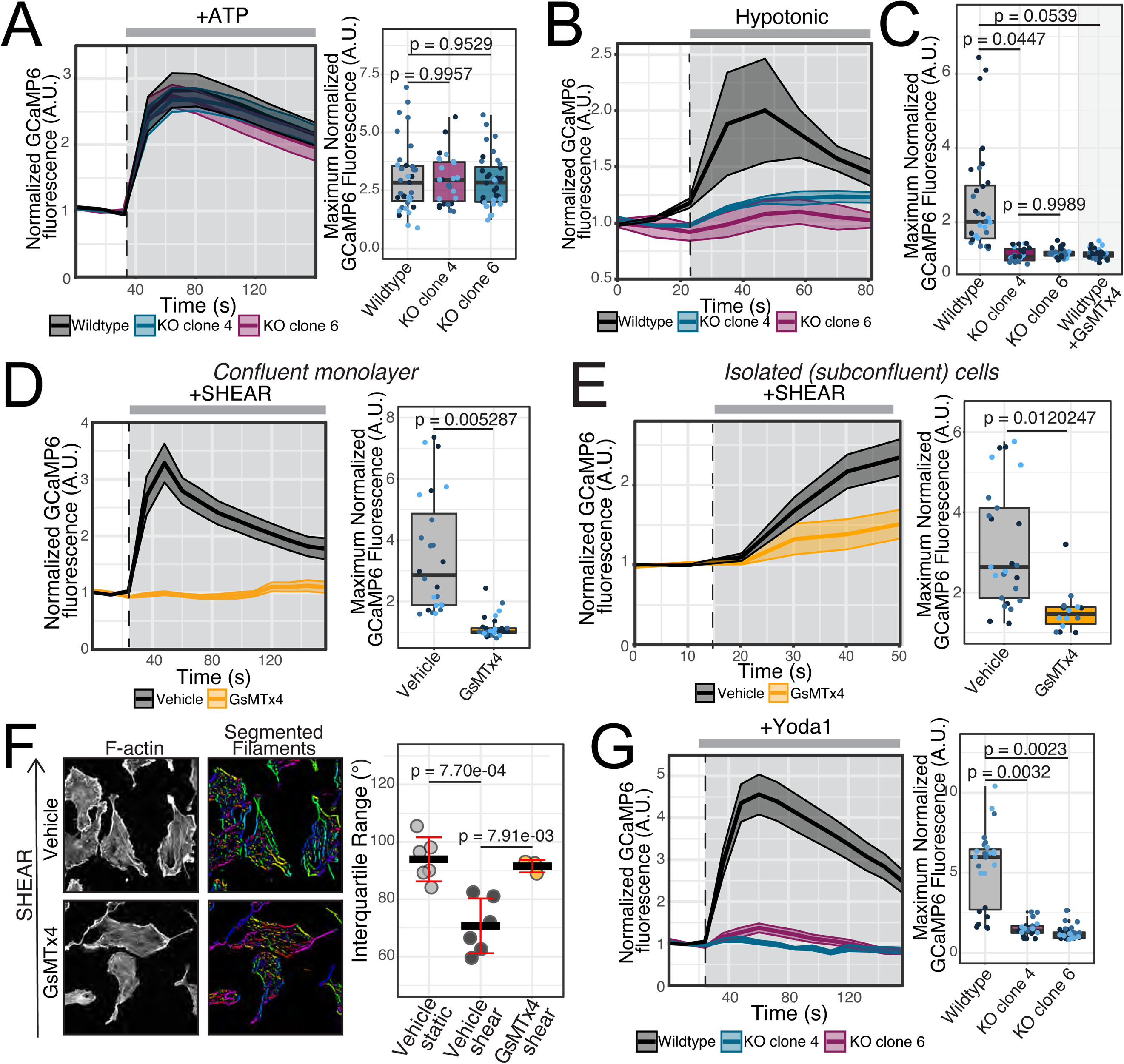
Spectrin-dependent shear-induced Ca^2+^ signaling is mediated by Piezo1. **A-E,G)** Live imaging of endothelial cells expressing the cytosolic [Ca^2+^] indicator GCaMP6 before and after the introduction of indicated stimuli (grey bar). Agonists were introduced at 24 sec and images were acquired every 12 sec. *Left panels:* Quantification of mean GCaMP6 fluorescence normalized to initial resting value. *Right panels:* Maximum normalized GCaMP6 fluorescence evoked by shear. Data are means ± SE of 3 independent experiments, each quantifying ≥8 cells. **A)** Wild-type and spectrin-KO cells were treated with 50 µM ATP in static conditions. **B-C)** Wild-type and spectrin-KO cells were subjected to 150 mOsm hypotonic shock. Where indicated the cells were incubated with the small peptide GsMTx4 for 30 min prior to hypotonic stress. **D-E)** Wild-type cells in confluent monolayers (**D**) or subconfluent cultures of sparsely distributed cells (**E**) were treated with vehicle (PBS) or incubated with GsMTx4 for 30 min prior to application of shear. **F)** Subconfluent cultures of sparsely distributed cells were treated with vehicle (PBS, top) or GsMTx4 (bottom) for 30 min prior to exposure to flow for a further 30 min as in Fig. 1A. *Left:* F-actin staining; *middle:* color labelling according to orientation of the maximum Feret diameter of the segmented filament relative to the shear axis (0°). *Right:* interquartile ranges of the distribution of segmented filament orientations for the indicated conditions. **G)** Wild-type and spectrin-KO cells were treated with 2.5 mM Yoda1 where indicated (grey background) while in static conditions.

Notably, however, responses to other mechanical stimuli were also impaired in *β*II-spectrin-KO cells (Fig. 4B, 4C). Increasing plasma membrane tension by inducing osmotic swelling through hypotonic stress resulted in a significant increase in [Ca^2+^]_cyto_ in wild-type but not in *β*II-spectrin-KO cells (Fig. 4B, 4C). Importantly, swelling-induced [Ca^2+^]_cyto_ was regulated by stretch-activated ion channels since pre-treatment with the small peptide GsMTx4, an amphipathic peptide that inhibits cationic mechanosensitive currents (Bae et al., 2011), eliminated this response (S. Fig. 3 and Fig. 4C). This strongly suggested that the responsiveness of mechanosensitive ion channels was impaired in *β*II-spectrin-KO cells.

It has been demonstrated that the stretch-activated, non-selective cation channel Piezo1 mediates (at least part of) the shear-induced Ca^2+^ increases in endothelial monolayers and their alignment in the direction of flow (Li et al., 2014). Consistent with this report, pre-treatment of tHAEC cells with GsMTx4, which blocks Piezo1, was sufficient to abrogate the shear-evoked [Ca^2+^] elevation in confluent monolayers (Fig. 4D). Importantly, this treatment also blunted the Ca^2+^ responses normally seen in single cells (Fig. 4E). GsMTx4 also inhibited the alignment of sparsely distributed endothelial cells in response to shear (Fig. 4F). These observations further indicated that the indispensable role of mechanosensitive ion channels in endothelial responses to fluid flow is regulated independently of junctional complexes.

The involvement of the Piezo1 channel was then probed more precisely using the highly specific, small molecule agonist, Yoda1. In static culture, treatment of wild-type cells with Yoda1 induced increases in [Ca^2+^]_cyto_ that strongly resembled the dynamics observed in response to flow (compare Fig. 4G to 3G). In contrast, Yoda1 did not elicit any significant [Ca^2+^]_cyto_ changes in *β*II-spectrin-KO cells (Fig. 4G). From this, we concluded that the apical *β*II-spectrin network regulates the activity of Piezo1.

### Plasma membrane expression of Piezo1 is not regulated by spectrin

The unresponsiveness of *β*II-spectrin-KO cells to Yoda1 suggested two possibilities: either (1) spectrins are required for the plasma membrane retention of Piezo1; or (2) Piezo1 activity at the cell surface is somehow regulated by the spectrin cytoskeleton. To test the first possibility, we examined the expression and cellular distribution of Piezo1. Immunoblotting experiments revealed comparable concentrations of Piezo1 protein in wild-type and *β*II-spectrin-KO cells (Fig. 5A and see S. Fig. 4 for quantitation). By immunofluorescence, we observed that in wild-type cells, Piezo1 appeared as punctate structures that concentrated on the apical surface of the endothelial cells, consistent with its shear-sensing functions (Fig. 5B). This distribution was largely preserved in *β*II-spectrin-KO cells, where numerous plasmalemma-localized punctate structures were observed (Fig. 5B). It is therefore unlikely that the alterations in Piezo1 responsiveness reported in spectrin-deficient cells are attributable to changes in the expression or localization of the channel.

**Figure 5:**
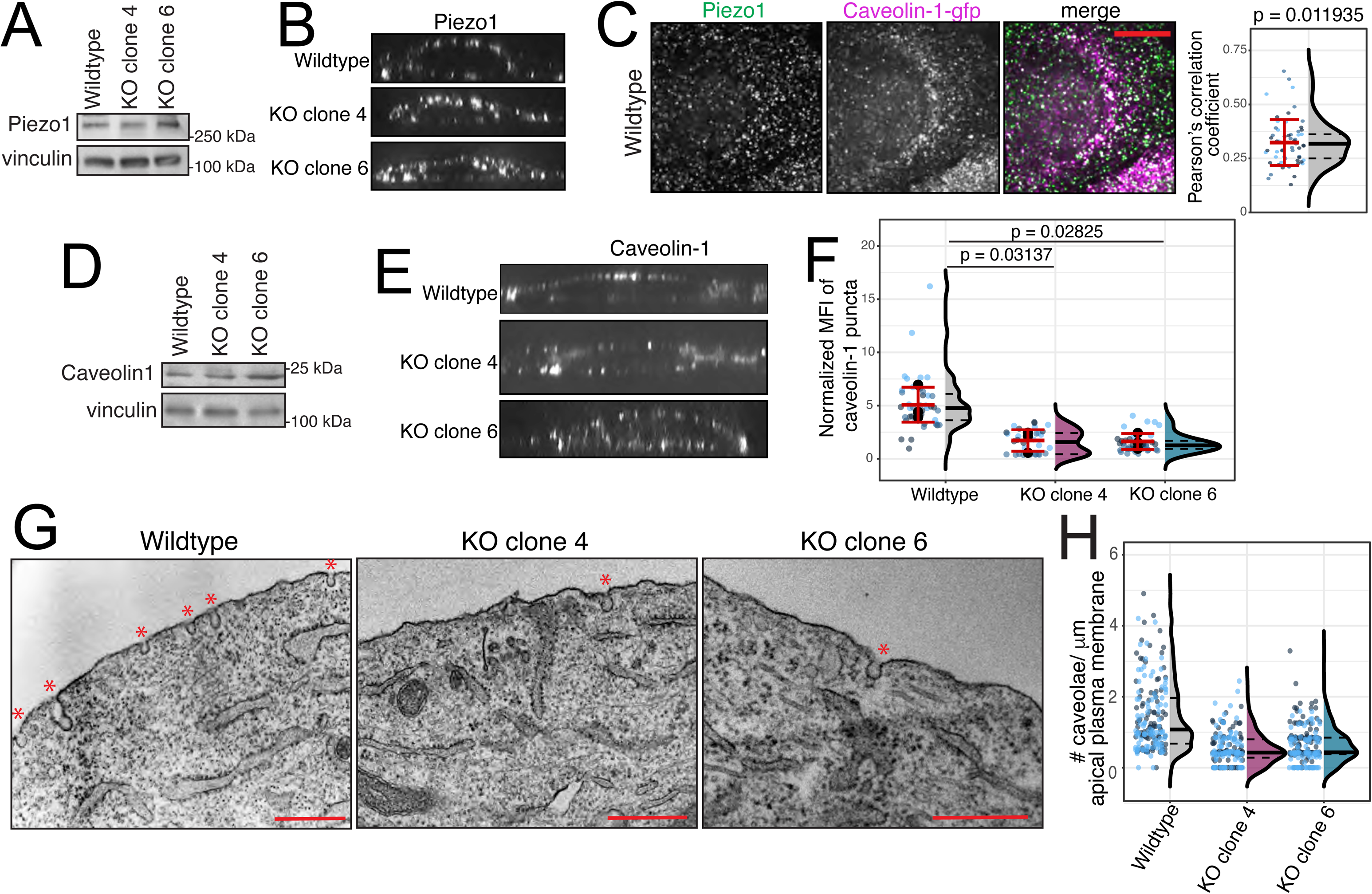
Piezo1 channels colocalize with caveolae which are stabilized by the spectrin cytoskeleton. **A)** Immunoblot of total Piezo1 expression in wild-type and spectrin KO cells; representative of 3 experiments. Vinculin was probed for normalization. **B)** Orthogonal confocal sections of cells immunostained for Piezo1; representative of 3 independent experiments. **C)** Cells expressing caveolin-GFP were subjected to methanol fixation and immunostained for Piezo1 (left, green in merge) and GFP (middle; magenta in merge). Representative confocal section from 3 independent experiments. Right: Pearson’s coefficient assessing the correlation between Piezo1 and caveolin, calculated for multiple individual cells. Jittered data points (in shades of blue) represent individual measurements and are color-coded according to biological replicate. Means of individual experiments are presented in black. Overall mean ± SE are indicated in red. Histogram of all data points indicating the mean (solid line) as well as the 25^th^ and 75^th^ percentiles (dashed line) are shown to the right of the individual data. **D)** Immunoblot of total caveolin-1 expression in wild-type and spectrin-KO cells; representative of 3 experiments. **E)** Orthogonal confocal sections of cells immunostained for caveolin-1; representative of 3 independent experiments. **F**) Analysis of the relative number of caveolin molecules in each caveolar cluster was performed by comparing the mean fluorescence of cellular puncta in immunostained cells to that of fluorescently conjugated secondary antibodies monodispersed on glass. Data are means ± SE of 3 independent experiments, each quantifying ≥20 cells. **G-H)** Sagittal sections of wild-type and spectrin-KO cells were analyzed by transmission electron microscopy. **G)** Representative electron micrographs of wildtype or spectrin KO endothelial cells with caveolae indicated by superimposed red asterisks. Scale bar = 500 nm. **H)** Number of caveolae per µm of plasma membrane in wild-type and spectrin-KO cells. From 2 independent experiments, each quantifying 90-100 cells.

### Spectrin stabilizes endothelial caveolae which regulate Piezo1

We proceeded to study how the spectrin-based membrane skeleton might directly impact Piezo1 function. Piezo1 expression is associated with local distortion of the lipid bilayer, often resulting in the preferential localization of the channels to curved membrane domains (Buyan et al., 2019; Diem et al., 2020; Liang and Howard, 2018). Localization to these domains is thought to confer a high degree of sensitivity to the channel, which responds to changes in membrane curvature that result from alterations in membrane tension (Diem et al., 2020; Liang and Howard, 2018).

Indeed, endothelial cells are uniquely enriched in curved membrane microdomains termed caveolae (Parton and Simons, 2007; Simionescu et al., 1981). Caveolae unfurl or flatten in response to application of force (Sinha et al., 2011). Of note, localization of Piezo1 to caveolar structures has previously been reported in other cells (Diem et al., 2020; Ridone et al., 2020). Accordingly, in wild-type endothelial cells, immunofluorescence revealed significant colocalization of Piezo1 and caveolin-1, a primary component of caveolae (Fig. 5C). We therefore investigated the effects of spectrin on caveolae, since these might indirectly influence Piezo1 activity.

Similar concentrations of total caveolin-1 protein were detected in wild-type and *β*II-spectrin-KO endothelial cells (Fig. 5D and see S. Fig. 4 for quantitation). When immunostained in wild-type endothelial cells, caveolin-1 formed punctate arrays that were enriched on the apical surface, consistent with the reported abundance of caveolae (Fig. 5E, F) (Drab et al., 2001; Oh et al., 2007). This polarized distribution was lost in *β*II-spectrin-KO cells, where a greater fraction of caveolin-1 was found in the basal and internal membranes (Fig. 5E). By analyzing the mean fluorescence intensities of the conjugated secondary antibodies within individual puncta on the apical cell surface, the number of caveolin-1 monomers in each cluster was also found to be significantly decreased in *β*II-spectrin-KO cells (Fig. 5F).

In addition to the observed changes in caveolin-1 distribution and clustering, we had previously observed that spectrin regulates the mobility of caveolar components in the plane of the membrane (Mylvaganam et al., 2020). However, it remained unclear whether the behavior of caveolin-1 reflected the number or appearance of the caveolae themselves. This was addressed by performing transmission electron microscopy (TEM) and analyzing sagittal sections of endothelial cells. In wild-type cells, TEM revealed a high apical density of uncoated, bulb-shaped plasmalemmal structures which we identified as caveolae (Fig. 5G, 5H). While still present, markedly fewer caveolae were observed on the surface of *β*II-spectrin-KO cells (Fig. 5G, 5H).

### Altered membrane tension in spectrin-deficient cells

Since caveolae play an important role in calibrating plasma membrane tension (Sinha et al., 2011), and given the relationship between spectrin expression and caveolin density, we wondered whether the resting membrane tension was altered in *β*II-spectrin-KO cells. To assess membrane tension, we performed fluorescence lifetime imaging microscopy (FLIM) using Flipper-TR, a membrane tension probe. The fluorescence lifetime of Flipper-TR increases at higher membrane tensions (Colom et al., 2018), which we confirmed by inducing osmotic swelling of wild-type cells with hypotonic (150 mOsm) medium (Fig. 6A). In isotonic conditions at rest, the fluorescence lifetime of flipper-TR in the plasma membrane of *β*II-spectrin-KO cells was significantly longer than the lifetime measured in their wild-type counterparts (Fig. 6B). This revealed that as in erythrocytes, spectrins maintain the flexibility of the endothelial plasma membrane. Elevated surface tension at rest in *β*II-spectrin-KO cells could account for the disappearance of caveolae, causing their flattening, and/or for a loss of the control of membrane tension afforded by caveolar structures. It could also account for the unresponsiveness of Piezo1 channels, which are known to desensitize when subjected to sustained increases in resting tension (Lewis and Grandl, 2015).

**Figure 6:**
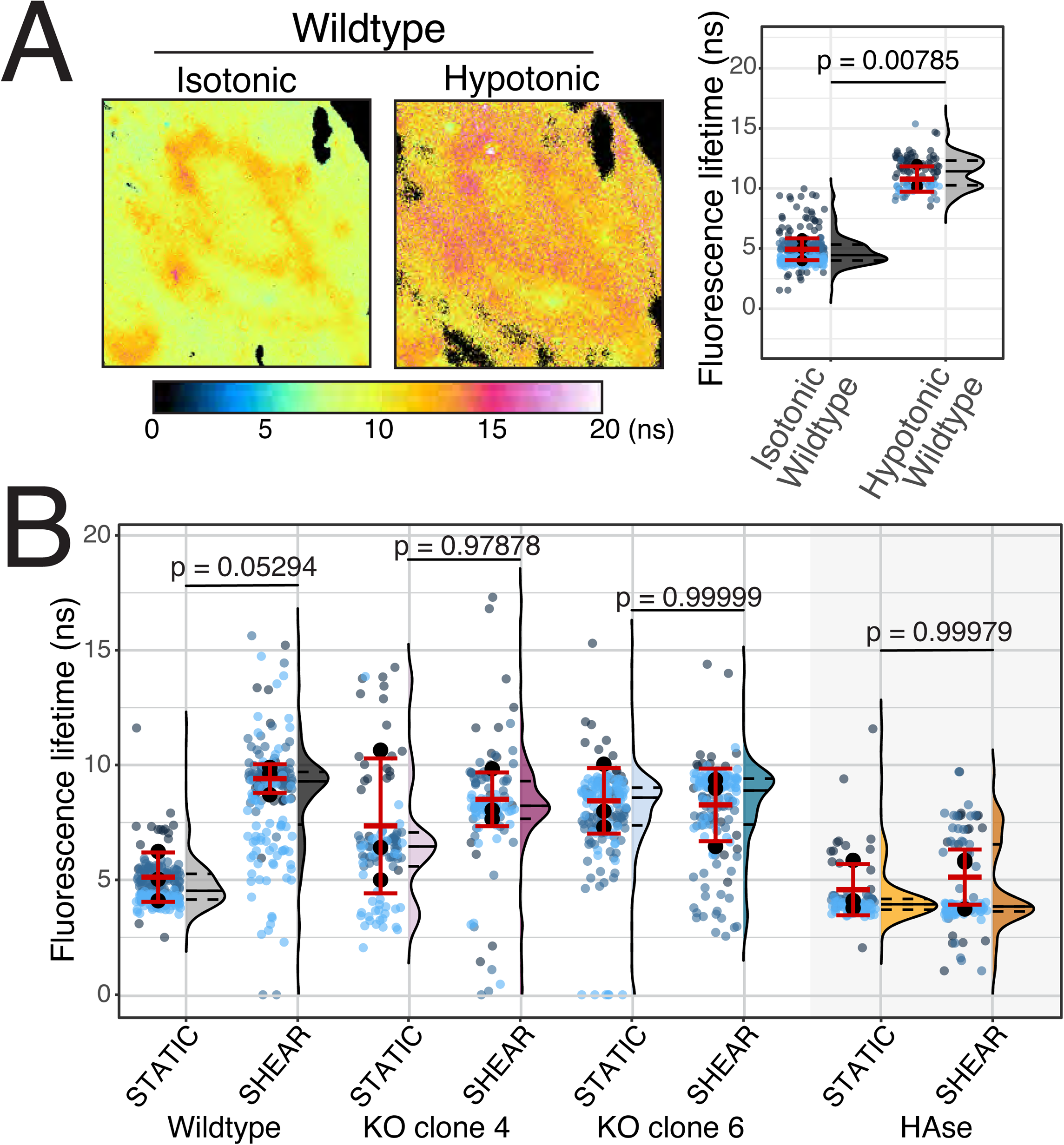
Effects of spectrin on plasma membrane tension. Cells were incubated with 1 µM Flipper-TR for 15 min and imaged by frequency-domain FLIM. **A)** Wildtype cells in static culture were measured before (*left*) and after (*middle*) exposure to 150 mOsm hypotonic shock. Representative heat-maps of fluorescence lifetimes (color bar: lifetime in nanoseconds (ns)) are illustrated. *Right panel*: mean lifetime of Flipper-TR estimated by frequency-domain FLIM under the indicated conditions. Here and in **B** jittered data points (in shades of blue) represent individual measurements and are color-coded according to biological replicate. Means of individual experiments are presented in black. Overall mean ± SE are indicated in red. Histogram of all data points indicating the mean (solid line) as well as the 25^th^ and 75^th^ percentiles (dashed line) are shown to the right of the individual data. Data from 3 independent experiments, each measuring ≥25 cells. **B)** FLIM measurements were performed prior to or 30 sec after introduction of shear stress. Mean lifetime of Flipper-TR in the specified cells is indicated. Data from 3 independent experiments, each measuring ≥25 cells.

### Spectrin and HA regulate shear-induced changes in membrane tension

We then studied the effects of shear on membrane tension. Shear stress acutely induced a significant increase in Flipper-TR fluorescence lifetime in wild-type cells, revealing an increase in plasma membrane tension (Fig. 6B). These changes were observed across the entire apical cell surface and did not appear to localize preferentially to any specific regions. In spectrin-KO cells, while shear induced an increase in Flipper-TR fluorescence lifetime (Fig. 6B), the magnitude of this increase was markedly lower than that observed in wild-type cells (Fig. 6B). Conceivably, this more modest increase in membrane tension was insufficient to activate Piezo1, which may already be rendered less sensitive/desensitized due to the higher resting tension in these cells.

While clearly important, spectrin is unlikely to directly sense the apical shear flow in order to increase membrane tension. Given its high extracellular density and established role in mediating shear-induced Ca^2+^ signaling events, it seemed more conceivable that HA, via its receptors, might instead transmit forces from the vascular lumen to the spectrin cytoskeleton to then induce and distribute changes in tension. Indeed, while treatment with HAase had no appreciable effect on membrane tension of cells in static culture, it greatly inhibited the shear-induced increase in the lifetime of Flipper-TR (Fig. 6B).

### Shear induced production of nitric oxide is regulated by spectrins

The preceding experiments indicated that spectrin behaves as a central integrator for endothelial signaling responses to flow. We therefore decided to further examine the importance of the cytoskeletal protein in general vascular functions. As mentioned, shear-induced production of NO is essential to maintain vascular health; NO both regulates changes in vascular tone and has important anti-coagulant properties. In the endothelium, NO is primarily produced by endothelial nitric oxide synthase (eNOS) which under basal conditions, interacts with caveolin-1. Caveolin-1 maintains the enzyme in an inactive state by sequestering it within caveolae. Increases in [Ca^2+^]_cyto_ result in the activation of calmodulin which, in its Ca^2+^-complexed form, can bind eNOS, disrupting its inhibitory interaction with caveolin-1 (Michel et al., 1997). Once dissociated from caveolae, eNOS can be activated through phosphorylation by multiple kinases (Bir et al., 2012; Calvert et al., 2008; Dimmeler et al., 1999; Fleming et al., 2001). Given that spectrin regulates both shear-induced [Ca^2+^]_cyto_ increases and the stability of caveolin-1/caveolae, it was likely that NO production might also be regulated by this network.

This was tested by immunoblotting cell lysates with specific antibodies that detect the phosphorylation of Ser1177 of eNOS. As shown in Fig. 7A, while application of shear (applied by vigorous rotary agitation) induced a distinct increase in eNOS phosphorylation in wild-type cells, no significant phosphorylation changes were recorded in *β*II-spectrin-KO cells.

**Figure 7:**
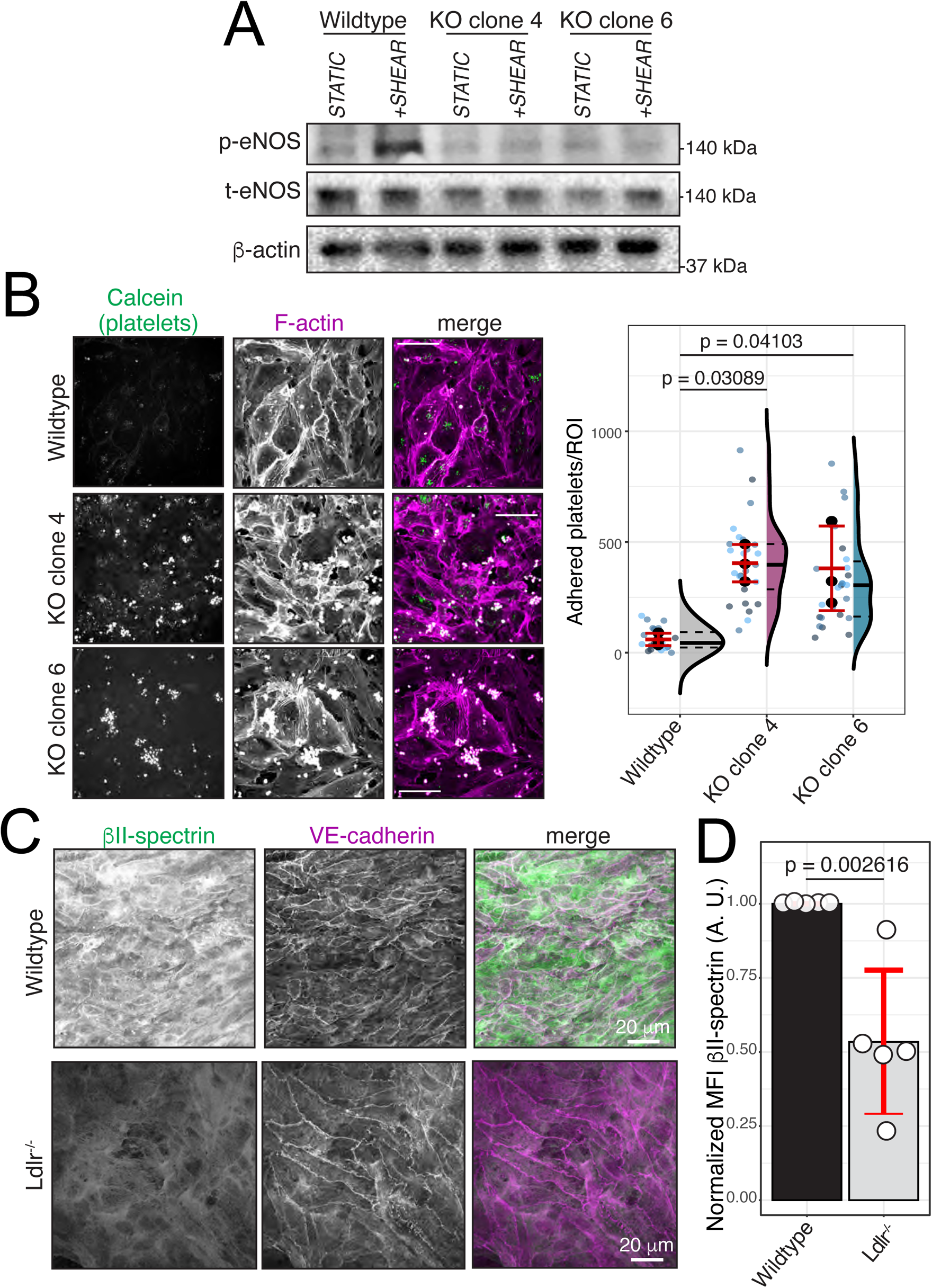
Association between spectrin deficiency and endothelial dysfunction. **A)** Representative immunoblot comparing eNOS phosphorylated on S1173 (p-eNOS;) and total eNOS (t-eNOS) in wild-type and spectrin-KO cells grown in static culture or following 10 min of shear stress applied by vigorous agitation on a laboratory shaker. **B)** The indicated endothelial cells lines were grown in collagen-coated microfluidic channels. Adhesion of calcein-labelled platelets was assessed under a constant shear flow rate of 15 dyne/cm^2^. *Left:* representative images. *Right:* platelet adhesion events scored per region of interest (ROI) analyzed for each condition after 5 min perfusion. Data from 3 independent experiments for each condition. **C-D)** *En face* aortic samples from wild-type (C57BL/6J) and *Ldlr*^−/−^ mice on a high fat diet for 4 weeks were prepared as described in Methods. **C)** Representative immunofluorescence images of *β*II-spectrin and VE-cadherin in wild-type and *Ldlr^−/−^* mouse aortas. **D)** Normalized mean fluorescence intensity of *β*II-spectrin, assessed by immunofluorescence. Data are means ± SE of 5 aortas from wild-type and *Ldlr^−/−^* mice, from two independent experiments.

To examine whether the impaired phosphorylation of eNOS translated into a functionally significant reduction in NO production, we examined the interaction between platelets and endothelial cells under flow. Very few platelets adhered to wild-type endothelial cells, consistent with the ability of NO to inhibit the attachment of platelets to the vessel wall. However, the number of platelets that adhered firmly to *β*II-spectrin-KO endothelial monolayers increased markedly, primarily along endothelial cell borders (Fig. 7B). These findings revealed that spectrin contributes to the anti-thrombotic properties of the endothelium.

We wondered whether changes in spectrin expression are associated with endothelial dysfunction in certain pathological contexts. Previous studies revealed a relationship between decreased expression of spectrins and atherosclerotic plaque severity (Rademakers et al., 2018). Increased activity of the protease calpain, which cleaves spectrin subunits leading to network disassembly, is also associated with early stages of atherosclerosis (Miyazaki et al., 2011). However, it remains unclear whether these changes in spectrin expression were reflective of the endothelium or other cells in the artery. To investigate this possibility, we examined the endothelial expression of *β*II-spectrin in *en face* preparations of descending thoracic aortas. We immunostained aortas of wild-type and *Ldlr^−/−^* mice that had been placed on a high-fat diet for four weeks to promote atheroma formation, according to previously established protocols. Consistent with previous observations, in contrast to their wild-type counterparts, diseased descending aortas were associated with a loss of endothelial cell alignment (Chien, 2007; Hahn and Schwartz, 2009). Interestingly, the aortic endothelium of wild-type mice expressed high levels of *β*II-spectrin (Fig. 7C-D). However, in *Ldlr^−/−^* aortas where plaques had formed, loss of alignment was coincident with a significantly lowered expression of *β*II-spectrin (Fig. 7C-D). These observations are generally consistent with a role of spectrin in promoting the normal atheroprotective functions of the endothelium and implicate its dysregulation in the sequelae of atherosclerosis.

## Discussion

While it has long been appreciated that the endothelium plays a central role in regulating vascular responses to shear stress, a comprehensive understanding of how these cells sense and transduce changes in blood flow has remained elusive. Here, we revisited the contribution of multiple proposed mechanotransducers to the shear-sensing functions of the endothelium and investigated mechanisms coordinating their activities. Importantly, we confirm that the junctional mechanosensory complex, which is central to current models of endothelial mechanotransduction, is not strictly required for endothelial responses to fluid flow (Fig. 1A, 1H). Other molecules were therefore likely to play an important role in directing these functions. We focused specifically on the functions of HA and Piezo1, which are indispensable for shear-induced responses of the endothelium.

### The spectrin cytoskeleton in mechanotransduction

It has been proposed that the cytoskeleton provides an integrative mechanism to distribute forces from mechano-sensors to mechano-responsive molecular complexes in cells (Davies, 1995). However, at least in the endothelium, very little experimental information exists for how this might occur. The majority of studies have focused on the actin cytoskeleton, largely because it plays a fundamental role in regulating the activity of integrins and junctional protein complexes that are often associated with mechanotransduction (Hahn and Schwartz, 2009). However, actin filaments cannot withstand large strains and are generally highly dynamic (Palmer et al., 1999). This would suggest that actin-rich structures alone are not well-suited to provide a framework for mechanosensitive cells.

We instead identified the spectrin cytoskeleton, which contains only short, relatively stable filaments of actin, as a fundamental integrator of mechanical signaling on the apical surface of the endothelium. The contributions of these networks to non-erythroid, non-neuronal cellular responses are rather poorly understood, perhaps due to the lack of pharmacological agents that target spectrin selectively. However, spectrin expression is highly conserved in metazoan tissues (Bennett and Healy, 2009). Furthermore, spectrins are consistently found to stabilize cell membranes and modulate the activity of some ion channels (Bennett and Healy, 2009). Though the underlying mechanism remains obscure, spectrins seem especially important to vascular function, as global deletion of *α*II- or *β*II-spectrin, the dominant isoforms expressed in non-erythroid cells, is lethal during embryonic development in mice (Stankewich et al., 2011) and zebrafish (Voas et al., 2007). Of note, deletion of *β*II-spectrin was associated with abnormal cardiac development at mid-gestation (Stankewich et al., 2011).

Spectrin proteins are extremely flexible and associate with short actin filaments that become highly stable; the resulting spectrin-actin lattice is uniquely pliable and long-lived (Lambert and Bennett, 1993). Flexibility of the cortical cytoskeleton confers increased compliance/deformability to the cell membrane and is therefore ideal for cells exposed to the dynamic environment of the vasculature. Indeed, the spectrin cytoskeleton is required to maintain the integrity of erythrocytes that experience hemodynamic stresses (Sheetz et al., 1980; Sheetz and Singer, 1977) and must squeeze through narrow capillaries (Agre et al., 1982; Chasis et al., 1988; Lux et al., 1976). Endothelial cells are similarly subjected to hemodynamic stress, especially in the aorta, and in narrow capillaries the endothelium is reciprocally deformed by passing erythrocytes. It is therefore unsurprising that spectrin plays a central role in blood vessel homeostasis. Furthermore, the vasculature is not the only environment where cells regularly experience fluctuations in pressure and shear. Cells in the lung are subjected to cyclic fluctuations in pressure. Compressions and tensions that arise from load-bearing activities result in fluid shear, both in the synovium of joints and in the lacunae of bones. The potential role of spectrin in these contexts should be explored.

### Caveolae and regulation of membrane tension

Importantly, in the endothelium, we find that spectrin is responsible for the maintenance of caveolae at the plasma membrane (Fig. 5G, 5H). As an important source of membrane reserve, caveolae contribute to the deformability of cells (Sinha et al., 2011) and they have been proposed to be involved in shear sensing (Parton and Simons, 2007). However, it has been unclear if shear induces flattening of these curved microdomains, or whether caveolae serve primarily as organizers of mechanosensitive molecular complexes. A previous study reported that osmotically-induced swelling of endothelial cells resulted in the flattening or remodeling of ~30% of their caveolae (Sinha et al., 2011). In our experiments, a similar hypo-osmotic stress resulted in a roughly two-fold increase in the fluorescence lifetime of the plasma membrane tension sensor Flipper-TR (Fig. 6A). This increase was comparable to that induced by the application of shear stress (Fig. 6B). The similarity of the changes in membrane tension support the hypothesis that (a fraction of) caveolae flatten in response to shear. However, electron microscopy studies will be required to confirm whether this is the case.

Because fewer caveolae were observed in the plasma membrane of spectrin-deficient cells, we speculated that remodelling of the cortical cytoskeleton –perhaps by magnifying the contribution of longer actin filaments– increased the resting tension of the apical membrane (Fig. 5G, 5H). Accordingly, the fluorescence lifetime of Flipper-TR in *β*II-spectrin-KO cells was longer than that of wild-type cells. The higher resting tension of the membrane of KO cells precluded any further increases when shear was applied to them, likely accounting for their failure to respond to fluid flow with increased [Ca^2+^].

Our data establish a correlation between caveolar stability/retention and the activity of Piezo1. Localization of Piezo1 to curved microdomains is largely credited for its high mechano-sensitivity (Guo and MacKinnon, 2017) and other studies indicate that resting membrane tension tunes the sensitivity of these channels (Lewis and Grandl, 2015). We therefore posit that spectrin regulates the sensitivity and activation of Piezo1 in two ways: by influencing the density of caveolae, and by maintaining a comparatively low apical membrane tension. It is also noteworthy that functional Piezo channels are homotrimeric complexes (Saotome et al., 2018) and that proper trimerization is likely required to induce plasma membrane curvature. Given our previous observations that spectrin regulates the diffusion and clustering of proteins in the plasma membrane (Mylvaganam et al., 2020), it is tempting to consider that the spectrin-dependent cytoskeletal network may dictate the diffusion and oligomerization of Piezo1 protomers and/or of other proteins that multimerize to generate caveolae.

In the context of vascular responses, caveolae are also important regulators of eNOS, which in unstimulated endothelial cells is inhibited by its direct association with caveolin-1. Ca^2+^ signaling induces dissociation of eNOS from caveolin-1, thereby promoting its activation (Fleming, 2010; Michel et al., 1997; Shaul et al., 1996). In view of the effects of spectrin deletion on caveolar stability and on Piezo1 responsiveness, the observed failure of spectrin KO cells to generate NO in response to flow was anticipated.

### A model of spectrin cytoskeleton-dependent mechanosensing

Our observations provide a tentative unifying framework to explain how responses to shear are integrated and communicated to intracellular effectors in endothelial cells. We believe that components of the glycocalyx, notably HA, are the primary sensors of fluid shear and that the primary HA receptor, CD44, is essential to convey the information. Accordingly, CD44 is required for normal alignment of endothelial cells in response to shear *in vitro* (Fig. 2B).

Interestingly, others ruled out mechanosensitive functions of CD44 because it lacks catch-bonding character (Mehta et al., 2020). However, CD44 is firmly linked to the spectrin cytoskeleton, which controls its distribution and mobility (Mylvaganam et al., 2020). It is therefore natural to envisage transfer of mechanical strain from CD44 to the underpinning spectrin meshwork. The latter is also linked to caveolae, dictating their stability and distribution. Displacement or structural changes in the spectrin network could readily alter the curvature or strain applied to caveolae and hence activate Piezo1. The resulting calcium influx, perhaps along with the mechanical distortion of the caveolae, stimulates the release and activation of eNOS. In this model spectrin plays a dual role: 1) it regulates the stability of glycocalyx components (that most directly experience fluid flow) and can sense and distribute mechanical disturbances experienced by the glycocalyx, and 2) it modulates plasma membrane tension by stabilizing curved membrane microdomains such as caveolae to influence the activation of mechanosensitive ion channels.

## Acknowledgments

S.M. is supported by a Vanier Scholarship from the Canadian Institutes of Health Research (CIHR) and a SickKids Restracomp Studentship. S.A.F and S.G. are supported by grants PJT-169180 and FDN-143202 from CIHR.

## Author Contributions

SM conducted experiments and data analyses. BY induced hypercholesterolemia in mice and performed dissections. RL acquired electron micrographs and CL performed platelet isolations. LAS provided reagents. SM, SAF and SG designed the study and wrote the manuscript with input from all authors.

## Methods

### Cell culture and transfections

Primary human aortic endothelial cells (HAEC), primary human umbilical vein endothelial cells (HUVEC), immortalized human aortic endothelial cells (tHAEC) and immortalized human umbilical vein endothelial cells (RF24) were grown in endothelial growth medium (EGM-2) containing 5% heat-inactivated fetal calf serum (FCS) with supplements (endothelial cell growth supplement: epidermal growth factor, basic fibroblast growth factor, heparin, and hydrocortisone, from PromoCell). Primary human adipose microvascular endothelial cells (HAMEC) were grown in endothelial cell medium (ECM) containing 5% heat-inactivated fetal bovine serum (FBS) and endothelial cell growth supplements (ScienCell) as previously described (Jaldin-Fincati et al., 2018). Cell cultures were maintained at 37°C in a 5% CO_2_ incubator with 95% humidity. Transfections were performed using the Neon electroporation system (Invitrogen) according to the manufacturer’s recommended protocol.

### Mice

Wild-type (C57BL/6J) and *Ldlr*^−/−^ mice were bred and housed under pathogen-free conditions at The Hospital for Sick Children animal facility. *In vivo* experiment protocols (Animal User Protocol—1000043631) were reviewed and approved by the Animal Care Committee of The Hospital for Sick Children in accordance with guidelines established by the Canadian Council on Animal Care (CCAC) and federal and provincial legislation.

### Reagents and plasmids

Antibodies for immunostaining were used as follows: rabbit anti-human caveolin-1 (BD Biosciences) at 1 µg/mL, mouse anti-human *β*II-spectrin (Santa Cruz Biotechnology) at 2 µg/mL, mouse anti-phosphotyrosine (4G10^®^ Platinum, Millipore Sigma) at 1 µg/mL, rabbit anti-Piezo1 (Novus Biologicals) at 4.5 µg/mL, goat anti-VE-Cadherin (Novus Biological) at 1 µg/mL, rabbit anti-CD31/PECAM-1 (Novus Biologicals) at 1 µg/mL. Cy3-, Alexa 488- or Alexa 647-conjugated donkey anti-mouse, anti-goat, or anti-rabbit secondary antibodies (Jackson ImmunoResearch) were used at 1 µg/mL. Rhodamine- or Alexa 488-phalloidin (Life Technologies) were used at 1:1000. Coverslips were pre-coated with rat-tail collagen type I (Sigma) at 50 µg/mL in PBS at 25°C for 1 h. Antibodies for immunoblotting were used as follows: mouse anti-human *β*II-spectrin at 1:5000 dilution, anti-Piezo1 at 1:1000 and anti-caveolin1 at 1:5000. phospho-eNOS Serine 1177 (Cell Signalling Technology) at 1:2500 dilution, anti-pan eNOS (Cell Signalling Technology) at 1:1000 dilution.

Hyaluronidase from bovine testes (Sigma) was used at 20 units/mL for 10-30 min at 37°C in DMEM. The Quantikine ELISA Hyaluronan Immunoassay (R&D Systems) was used as per manufacturer’s instructions. Caveolin-1-RFP was previously described (Hirama et al., 2017). GCaMP6 was used as described (Chen et al., 2013). Stock solutions of Flipper-TR (Spirochrome) were prepared in DMSO and used according to manufacturer’s instructions. Mouse high-fat diet D12108C (Clinton/Cybulsky High-Fat Rodent Diet With Regular Casein and 1.25% Added Cholesterol) was used, supplied by Research Diets Inc.

### CRISPR/Cas9 and RNA-interference

*β*II-spectrin-KO CRISPR cell lines were generated as follows: a predesigned vector containing a human *β*II-spectrin-targeting gRNA and eCas9-2A-tGFP was purchased from Millipore Sigma (MISSION gRNA ID: HSP00000399630). 24 hours after transfection of this plasmid, single GFP-positive cells were sorted into 96-well plates and expanded for 2-3 weeks to generate clonal populations. *β*II-spectrin deletion was confirmed in clones by immunoblotting. Stealth RNAi^TM^ siRNAs (ThermoFisher) were electroporated at 100 pM using the Neon transfection system.

### Shear stress

Endothelial cell monolayers were grown in glass-bottom microfluidic channels (Ibidi) coated with 50 µg/mL rat tail collagen (Sigma). Inlets of the microfluidic channels were connected with tubing to syringes filled with complete media in a syringe pump (Harvard Apparatus), while outlet tubing was connected to a waste container. The syringe pump was used to apply a fluid flow rate of 3.9 mL/min which yields an estimated shear rate of 15 dyne/cm^2^ (Buchanan et al., 2014).

### Immunofluorescence

Endothelial cell monolayers were grown on coverslips or in microfluidic channels coated with 50 μg/mL rat tail collagen. Cells were fixed with 4% paraformaldehyde (PFA) (Electron Microscopy Sciences), permeabilized with 0.1% Triton X-100 in PBS, blocked with 2% BSA in PBS, and stained with the indicated antibodies or fluorescent phalloidin in blocking solution. After staining, coverslips were mounted onto glass slides using ProLong Diamond (Life Technologies) or imaged directly in PBS.

### Confocal microscopy and image analysis

Unless otherwise indicated, imaging was performed using a Quorum spinning disc system mounted on a Zeiss Axiovert 200M microscope, using 10x air, 25x water, 63x oil or 100x oil objectives, a 1.5x magnification lens, and a back-thinned EM-CCD camera (C9100-13, Hamamatsu). Acquisitions were controlled by the Volocity software (Perkin-Elmer), exported and processed with MatLab (MathWorks) for single-particle tracking and filament alignment analysis, or analyzed and quantified using Volocity or FIJI/Image J (NIH) software.

### Analysis of F-actin orientations

Confocal images of F-actin were analyzed using custom MATLAB scripts using MATLAB_R2020a. Briefly, images were skeletonized using a Hessian-based multiscale filter and converted to a binary label matrix. Tubular structures, or “filaments,” were then segmented by fitting structural elements to the skeleton. Segmented areas smaller than 3 pixels were excluded. The maximum Feret diameter of each segmented filament was identified as the longest possible distance between two parallel tangents at opposing borders of the structural element. The angle between the maximum Feret diameter relative to the direction of applied flow was then calculated.

### Transmission electron microscopy (TEM) and analysis

TEM was performed according to standard methods. Briefly, tHAEC were fixed in a mixture of 2% PFA, 2.5% glutaraldehyde and 0.1 M cacodylate buffer. Subsequently, the cells were post-fixed with 1% osmium tetroxide, dehydrated in a graded series of ethanol before embedding in Quetol-Spurr resin. Thin sagittal sections were cut and stained with uranyl acetate and lead citrate. Grids were examined using a JEOLJEM-1011 electron microscope at 80 kV with a side-mounted Advantage HR CCD camera (Advanced Microscopy Techniques).

### Fluorescence lifetime imaging microscopy (FLIM)

Cells were incubated with 1 µM flipper-TR for 15 min prior to imaging. Frequency-domain FLIM measurements were collected using an Olympus IX81 inverted microscope equipped with a Lambert-FLIM attachment, using a 60x/1.49 NA oil immersion objective and Li2CAM iCCD camera. The frequency of modulation was 400 MHz and the instrument was calibrated based on the fluorescence lifetime of AlexaFluor 546 assuming a monoexponential lifetime of 4.1 ns (Berezin and Achilefu, 2010). Lifetime analysis was performed using Lambert Instruments FLIM software.

### Platelet isolation

Whole blood was collected from healthy donors using 0.6% Acid-Citrate-Dextrose solution A (ACD-A) as the anticoagulant. Washed platelets were prepared using a light-spin/hard-spin method and 0.6% ACD-A/PBS solution.

### Induction of hypercholesterolemia

Hypercholesterolemia was induced in 19-week-old wild-type (C57BL/6J) and *Ldlr*^−/−^ mice by replacing normal chow with high fat diet (1.25% cholesterol, D12108C, Research Diets Inc.) for 4 weeks.

### Aortic arch *en face* immunostaining

*En face* aortic samples were prepared as described previously (Paulson et al., 2010). Briefly, following intracardiac puncture, mice were perfused through the left ventricle with ice-cold PBS for 5 min followed by 4% PFA in PBS for 5 min. Once harvested, aortas were further fixed in 4% PFA for 24 hours at 4°C. Next, adipose tissue was dissected while immersed in cold PBS. Permeabilization was performed with 0.1% Triton-X in PBS for 30 min. Aortas were incubated overnight with primary antibodies at 4°C. The aorta was opened in a reproducible manner by cutting along the greater curvature then mounted on glass slides with mounting medium (Dako Fluorescent; DakoCytomation).

### Quantification and statistical analysis

The number of experiments and cells quantified are indicated in the individual Figure Legends. Statistical analysis was performed using R statistical software as indicated.

## Supplemental Figure Legends

**S. Fig 1:**
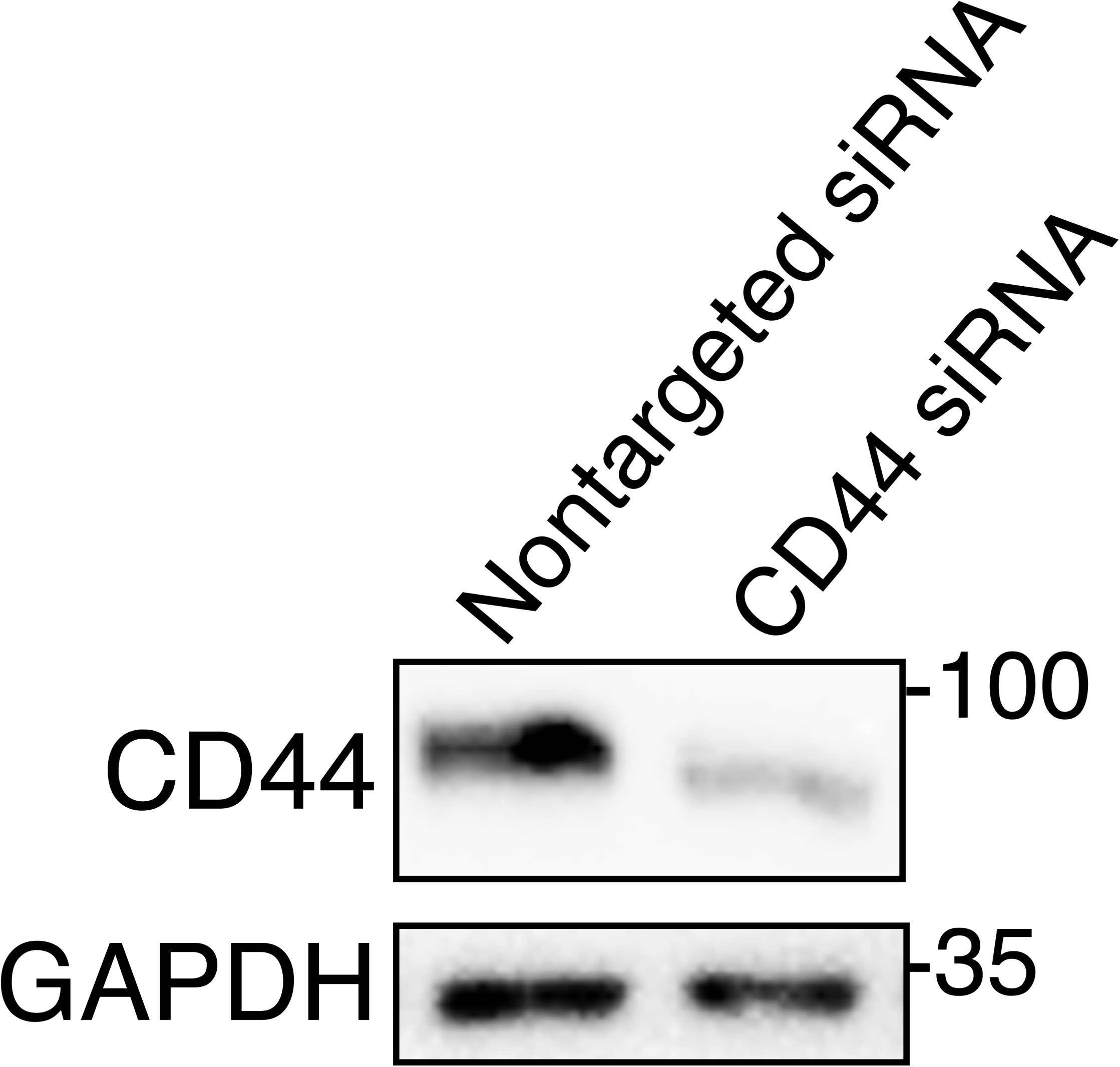
Silencing of CD44 expression in endothelial cells. CD44 expression in RF24 cells probed by immunoblotting after transfection with non-targeted or CD44-targeted siRNAs. Normalized to GAPDH.

**S. Fig 2:**
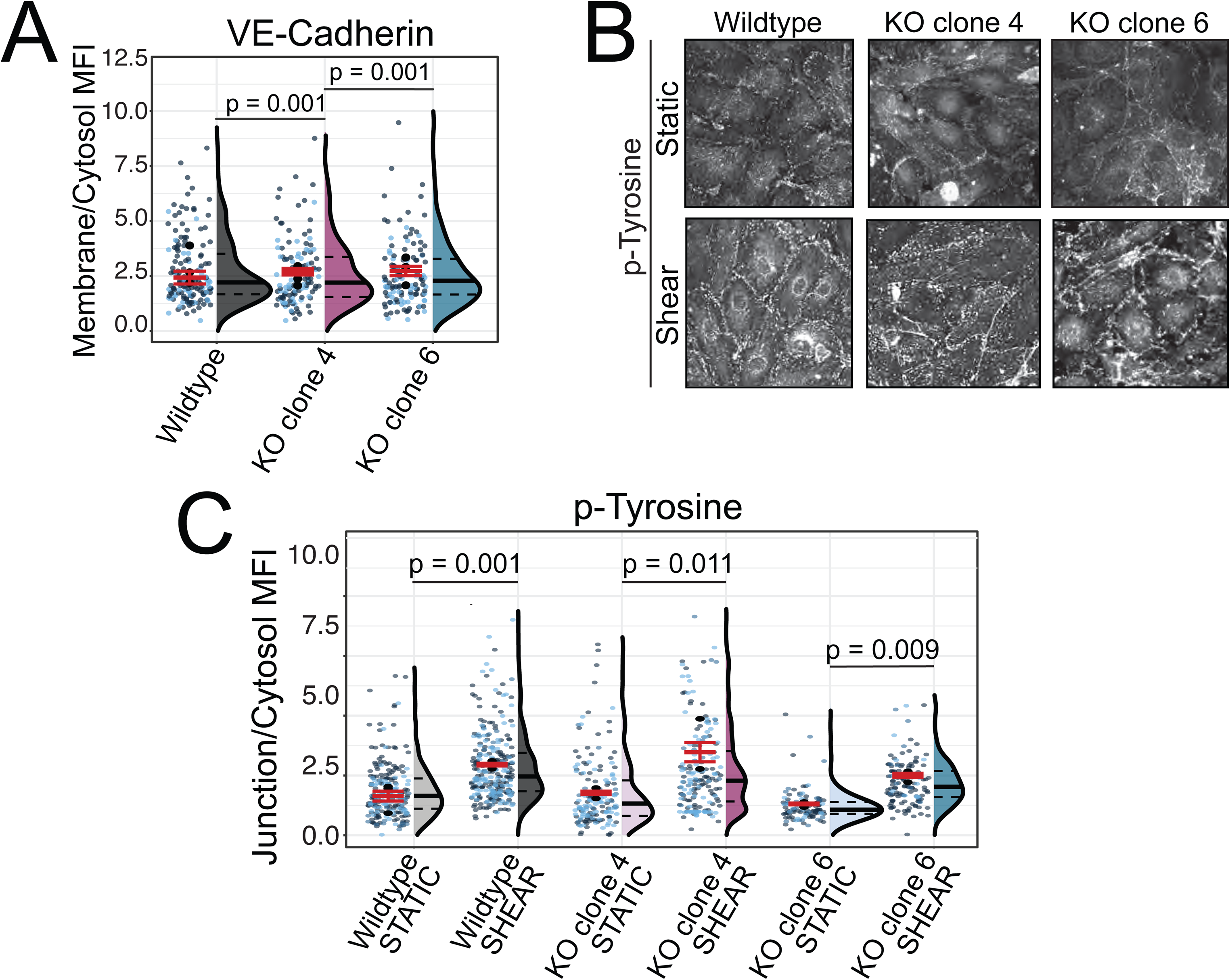
Shear-induced tyrosine phosphorylation of wildtype and βII-spectrin-KO cells. **A)** Ratio of VE-cadherin fluorescence at intercellular contacts and cytosol for individual cells in a confluent monolayer. In A and C jittered data points (in shades of blue) represent individual measurements and are color-coded according to biological replicate. Means of individual experiments are presented in black. Overall means ± SE are indicated in red. Histograms of all data points indicating the mean (solid line) as well as the 25^th^ and 75^th^ percentiles (dashed line) are shown to the right of the individual data. Data from 3 independent experiments, each quantifying ≥50 cells. **B-C)** Endothelial cells were subjected to constant fluid flow or maintained in static culture for 30 min. Cells were then fixed and immunostained for phosphorylated protein tyrosine residues (p-Tyrosine). **B)** Representative micrographs for indicated conditions. **C)** Ratio of phospho-tyrosine fluorescence at junctions (VE-cadherin-positive structures) and cytosol for individual cells. Data from 3 independent experiments, each quantifying ≥50 cells.

**S. Fig 3:**
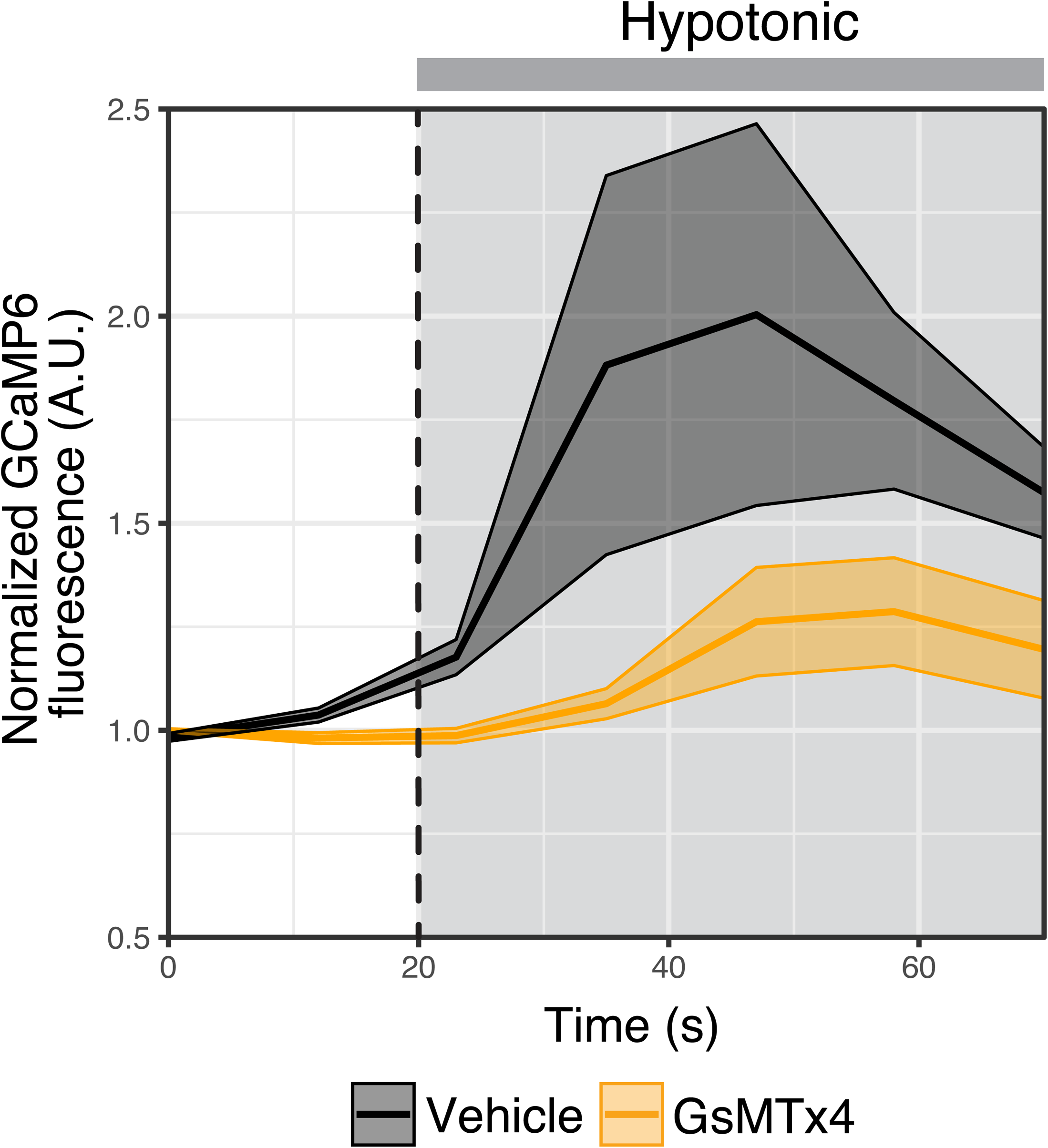
Effect of GsMTx4 on hypotonically-induced [Ca^2+^] changes. Live imaging of confluent endothelial cells expressing the cytosolic [Ca^2+^] indicator GCaMP6 before and after the introduction of hypotonic stress (grey background). Images were acquired every 12 sec. Cells were treated with vehicle (PBS) or the small peptide GsMTx4 for 30 min prior to imaging. Quantification of mean GCaMP6 fluorescence over time. Data are means ± SE (shaded area) of 3 independent experiments, each quantifying ≥8 cells

**S. Fig 4:**
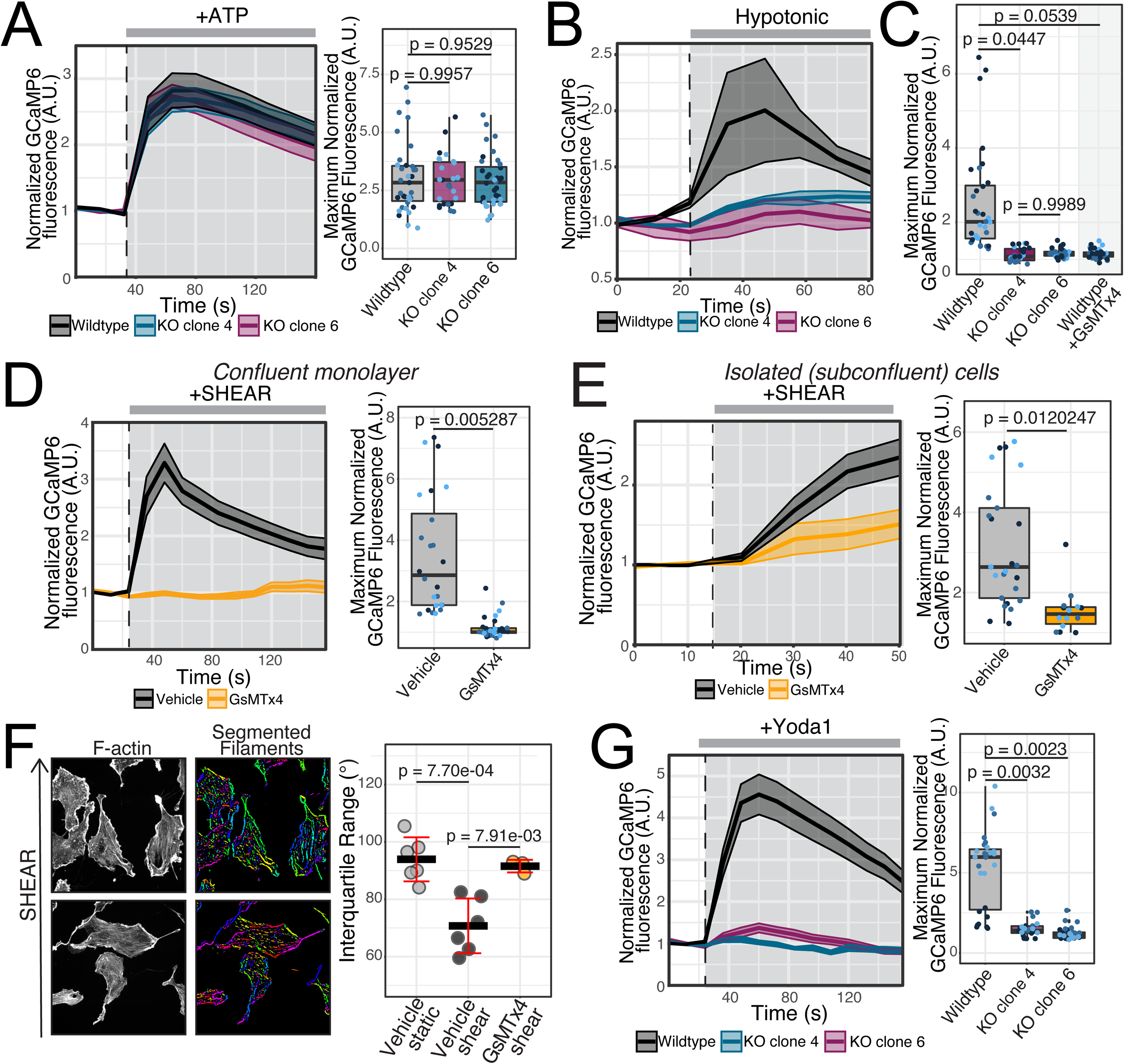
Comparable expression of Piezo1 and caveolin-1 proteins in wildtype and βII-spectrin-KO cells. Wild-type or spectrin-KO endothelial cells were lysed and (A) Piezo1 or (B) Caveolin-1 expression was assessed by immunoblotting and compared to vinculin as in Figure 5. Quantification of protein expression by densitometric analysis of 3 independent experiments, normalized in each instace to the wild-type control.

## Notes

### Competing Interest Statement

The authors have declared no competing interest.

## References

Agre, P., Bennett, V., and Orringer, E.P. (1982). Deficient red-cell spectrin in severe, recessively inherited spherocytosis.

Aruffo, A., Stamenkovic, I., Melnick, M., Underhill, C.B., and Seed, B. (1990). CD44 is the principal cell surface receptor for hyaluronate. Cell 61, 1303–1313.

Bae, C., Sachs, F., and Gottlieb, P.A. (2011). The mechanosensitive ion channel Piezo1 is inhibited by the peptide GsMTx4. Biochemistry 50, 6295–6300.

Baeyens, N., Bandyopadhyay, C., Coon, B.G., Yun, S., and Schwartz, M.A. (2016). Endothelial fluid shear stress sensing in vascular health and disease. J Clin Invest 126, 821–828.

Barbee, K.A., Davies, P.F., and Lal, R. (1994). Shear stress-induced reorganization of the surface topography of living endothelial cells imaged by atomic force microscopy. Circ Res 74, 163–171.

Barbee, K.A., Mundel, T., Lal, R., and Davies, P.F. (1995). Subcellular distribution of shear stress at the surface of flow-aligned and nonaligned endothelial monolayers. Am J Physiol 268, H1765–1772.

Bennett, V., and Baines, A.J. (2001). Spectrin and ankyrin-based pathways: metazoan inventions for integrating cells into tissues. Physiological reviews 81, 1353–1392.

Bennett, V., and Healy, J. (2009). Membrane domains based on ankyrin and spectrin associated with cell–cell interactions. Cold Spring Harbor perspectives in biology 1, a003012.

Berezin, M.Y., and Achilefu, S. (2010). Fluorescence lifetime measurements and biological imaging. Chem Rev 110, 2641–2684.

Bir, S.C., Xiong, Y., Kevil, C.G., and Luo, J. (2012). Emerging role of PKA/eNOS pathway in therapeutic angiogenesis for ischaemic tissue diseases. Cardiovasc Res 95, 7–18.

Buchanan, C.F., Voigt, E.E., Szot, C.S., Freeman, J.W., Vlachos, P.P., and Rylander, M.N. (2014). Three-dimensional microfluidic collagen hydrogels for investigating flow-mediated tumor-endothelial signaling and vascular organization. Tissue Engineering Part C: Methods 20, 64–75.

Buga, G.M., Gold, M.E., Fukuto, J.M., and Ignarro, L.J. (1991). Shear stress-induced release of nitric oxide from endothelial cells grown on beads. Hypertension 17, 187–193.

Buyan, A., Cox, C.D., Rae, J., Barnoud, J., Li, J., Cvetovska, J., Bastiani, M., Chan, H.S.M., Hodson, M.P., Martinac, B., et al. (2019). Piezo1 Induces Local Curvature in a Mammalian Membrane and Forms Specific Protein-Lipid Interactions. bioRxiv, 787531.

Calvert, J.W., Gundewar, S., Jha, S., Greer, J.J., Bestermann, W.H., Tian, R., and Lefer, D.J. (2008). Acute metformin therapy confers cardioprotection against myocardial infarction via AMPK-eNOS-mediated signaling. Diabetes 57, 696–705.

Camenisch, T.D., Spicer, A.P., Brehm-Gibson, T., Biesterfeldt, J., Augustine, M.L., Calabro, A., Kubalak, S., Klewer, S.E., and McDonald, J.A. (2000). Disruption of hyaluronan synthase-2 abrogates normal cardiac morphogenesis and hyaluronan-mediated transformation of epithelium to mesenchyme. The Journal of clinical investigation 106, 349–360.

Chasis, J., Agre, P., and Mohandas, N. (1988). Decreased membrane mechanical stability and in vivo loss of surface area reflect spectrin deficiencies in hereditary spherocytosis. The Journal of clinical investigation 82, 617–623.

Chen, T.-W., Wardill, T.J., Sun, Y., Pulver, S.R., Renninger, S.L., Baohan, A., Schreiter, E.R., Kerr, R.A., Orger, M.B., and Jayaraman, V. (2013). Ultrasensitive fluorescent proteins for imaging neuronal activity. Nature 499, 295–300.

Chien, S. (2007). Mechanotransduction and endothelial cell homeostasis: the wisdom of the cell. American Journal of Physiology-Heart and Circulatory Physiology 292, H1209–H1224.

Colom, A., Derivery, E., Soleimanpour, S., Tomba, C., Dal Molin, M., Sakai, N., González-Gaitán, M., Matile, S., and Roux, A. (2018). A fluorescent membrane tension probe. Nature chemistry 10, 1118–1125.

Conway, D.E., Breckenridge, M.T., Hinde, E., Gratton, E., Chen, C.S., and Schwartz, M.A. (2013). Fluid shear stress on endothelial cells modulates mechanical tension across VE-cadherin and PECAM-1. Curr Biol 23, 1024–1030.

Copp, A.J. (1995). Death before birth: clues from gene knockouts and mutations. Trends Genet 11, 87–93.

Davies, P.F. (1995). Flow-mediated endothelial mechanotransduction. Physiol Rev 75, 519–560.

Davies, P.F. (2009). Hemodynamic shear stress and the endothelium in cardiovascular pathophysiology. Nat Clin Pract Cardiovasc Med 6, 16–26.

Davies, P.F., Dewey, C.F., Jr., Bussolari, S.R., Gordon, E.J., and Gimbrone, M.A., Jr. (1984). Influence of hemodynamic forces on vascular endothelial function. In vitro studies of shear stress and pinocytosis in bovine aortic cells. J Clin Invest 73, 1121–1129.

Dewey, C.F., Jr., Bussolari, S.R., Gimbrone, M.A., Jr., and Davies, P.F. (1981). The dynamic response of vascular endothelial cells to fluid shear stress. J Biomech Eng 103, 177–185.

Diem, K., Fauler, M., Fois, G., Hellmann, A., Winokurow, N., Schumacher, S., Kranz, C., and Frick, M. (2020). Mechanical stretch activates piezo1 in caveolae of alveolar type I cells to trigger ATP release and paracrine stimulation of surfactant secretion from alveolar type II cells. The FASEB Journal 34, 12785–12804.

Dimmeler, S., Fleming, I., Fisslthaler, B., Hermann, C., Busse, R., and Zeiher, A.M. (1999). Activation of nitric oxide synthase in endothelial cells by Akt-dependent phosphorylation. Nature 399, 601–605.

Doucette, J.W., Corl, P.D., Payne, H.M., Flynn, A.E., Goto, M., Nassi, M., and Segal, J. (1992). Validation of a Doppler guide wire for intravascular measurement of coronary artery flow velocity. Circulation 85, 1899–1911.

Drab, M., Verkade, P., Elger, M., Kasper, M., Lohn, M., Lauterbach, B., Menne, J., Lindschau, C., Mende, F., Luft, F.C., et al. (2001). Loss of caveolae, vascular dysfunction, and pulmonary defects in caveolin-1 gene-disrupted mice. Science 293, 2449–2452.

Duncan, G.S., Andrew, D.P., Takimoto, H., Kaufman, S.A., Yoshida, H., Spellberg, J., De La Pompa, J.L., Elia, A., Wakeham, A., and Karan-Tamir, B. (1999). Genetic evidence for functional redundancy of platelet/endothelial cell adhesion molecule-1 (PECAM-1): CD31-deficient mice reveal PECAM-1-dependent and PECAM-1-independent functions. The Journal of Immunology 162, 3022–3030.

Fleming, I. (2010). Molecular mechanisms underlying the activation of eNOS. Pflugers Arch 459, 793–806.

Fleming, I., Fisslthaler, B., Dimmeler, S., Kemp, B.E., and Busse, R. (2001). Phosphorylation of Thr(495) regulates Ca(2+)/calmodulin-dependent endothelial nitric oxide synthase activity. Circ Res 88, E68–75.

Gimbrone, M.A., Jr., Topper, J.N., Nagel, T., Anderson, K.R., and Garcia-Cardena, G. (2000). Endothelial dysfunction, hemodynamic forces, and atherogenesis. Ann N Y Acad Sci 902, 230–239; discussion 239-240.

Guo, Y.R., and MacKinnon, R. (2017). Structure-based membrane dome mechanism for Piezo mechanosensitivity. Elife 6, e33660.

Hahn, C., and Schwartz, M.A. (2009). Mechanotransduction in vascular physiology and atherogenesis. Nature reviews Molecular cell biology 10, 53–62.

Hirama, T., Das, R., Yang, Y., Ferguson, C., Won, A., Yip, C.M., Kay, J.G., Grinstein, S., Parton, R.G., and Fairn, G.D. (2017). Phosphatidylserine dictates the assembly and dynamics of caveolae in the plasma membrane. J Biol Chem 292, 14292–14307.

Homan, K.A., Gupta, N., Kroll, K.T., Kolesky, D.B., Skylar-Scott, M., Miyoshi, T., Mau, D., Valerius, M.T., Ferrante, T., Bonventre, J.V., et al. (2019). Flow-enhanced vascularization and maturation of kidney organoids in vitro. Nat Methods 16, 255–262.

Hong, D., Jaron, D., Buerk, D.G., and Barbee, K.A. (2006). Heterogeneous response of microvascular endothelial cells to shear stress. American Journal of Physiology-Heart and Circulatory Physiology 290, H2498–H2508.

Jaldin-Fincati, J.R., Pereira, R.V.S., Bilan, P.J., and Klip, A. (2018). Insulin uptake and action in microvascular endothelial cells of lymphatic and blood origin. Am J Physiol Endocrinol Metab 315, E204–E217.

Jin, Z.-G., Ueba, H., Tanimoto, T., Lungu, A.O., Frame, M.D., and Berk, B.C. (2003). Ligand-independent activation of vascular endothelial growth factor receptor 2 by fluid shear stress regulates activation of endothelial nitric oxide synthase. Circulation research 93, 354–363.

Lambert, S., and Bennett, V. (1993). From anemia to cerebellar dysfunction: a review of the ankyrin gene family. European journal of biochemistry 211, 1–6.

Levick, J.R. (1991). An introduction to cardiovascular physiology (London; Boston: Butterworths).

Lewis, A.H., and Grandl, J. (2015). Mechanical sensitivity of Piezo1 ion channels can be tuned by cellular membrane tension. Elife 4, e12088.

Li, J., Hou, B., Tumova, S., Muraki, K., Bruns, A., Ludlow, M.J., Sedo, A., Hyman, A.J., McKeown, L., and Young, R.S. (2014). Piezo1 integration of vascular architecture with physiological force. Nature 515, 279–282.

Liang, X., and Howard, J. (2018). Structural biology: Piezo senses tension through curvature. Current Biology 28, R357–R359.

Lückhoff, A., Pohl, U., Mülsch, A., and Busse, R. (1988). Differential role of extra-and intracellular calcium in the release of EDRF and prostacyclin from cultured endothelial cells. British journal of pharmacology 95, 189–196.

Lux, S.E., JoHN, K.M., and Karnovsky, M. (1976). Irreversible deformation of the spectrin-actin lattice in irreversibly sickled cells. The Journal of clinical investigation 58, 955–963.

Mehta, V., Pang, K.-L., Rozbesky, D., Nather, K., Keen, A., Lachowski, D., Kong, Y., Karia, D., Ameismeier, M., and Huang, J. (2020). The guidance receptor plexin D1 is a mechanosensor in endothelial cells. Nature 578, 290–295.

Michel, J.B., Feron, O., Sacks, D., and Michel, T. (1997). Reciprocal regulation of endothelial nitric-oxide synthase by Ca2+-calmodulin and caveolin. J Biol Chem 272, 15583–15586.

Miyazaki, T., Taketomi, Y., Takimoto, M., Lei, X.F., Arita, S., Kim-Kaneyama, J.R., Arata, S., Ohata, H., Ota, H., Murakami, M., et al. (2011). m-Calpain induction in vascular endothelial cells on human and mouse atheromas and its roles in VE-cadherin disorganization and atherosclerosis. Circulation 124, 2522–2532.

Mochizuki, S., Vink, H., Hiramatsu, O., Kajita, T., Shigeto, F., Spaan, J.A., and Kajiya, F. (2003). Role of hyaluronic acid glycosaminoglycans in shear-induced endothelium-derived nitric oxide release. American Journal of Physiology-Heart and Circulatory Physiology.

Mylvaganam, S., Riedl, M., Vega, A., Collins, R.F., Jaqaman, K., Grinstein, S., and Freeman, S.A. (2020). Stabilization of Endothelial Receptor Arrays by a Polarized Spectrin Cytoskeleton Facilitates Rolling and Adhesion of Leukocytes. Cell Rep 31, 107798.

Necas, J., Bartosikova, L., Brauner, P., and Kolar, J. (2008). Hyaluronic acid (hyaluronan): a review. Veterinarni medicina 53, 397–411.

Newman, P.J. (1997). The biology of PECAM-1. The Journal of clinical investigation 99, 3–8.

Oh, P., Borgstrom, P., Witkiewicz, H., Li, Y., Borgstrom, B.J., Chrastina, A., Iwata, K., Zinn, K.R., Baldwin, R., Testa, J.E., et al. (2007). Live dynamic imaging of caveolae pumping targeted antibody rapidly and specifically across endothelium in the lung. Nat Biotechnol 25, 327–337.

Pahakis, M.Y., Kosky, J.R., Dull, R.O., and Tarbell, J.M. (2007). The role of endothelial glycocalyx components in mechanotransduction of fluid shear stress. Biochemical and biophysical research communications 355, 228–233.

Palmer, A., Mason, T.G., Xu, J., Kuo, S.C., and Wirtz, D. (1999). Diffusing wave spectroscopy microrheology of actin filament networks. Biophysical journal 76, 1063–1071.

Parton, R.G., and Simons, K. (2007). The multiple faces of caveolae. Nat Rev Mol Cell Biol 8, 185–194.

Paulson, K.E., Zhu, S.N., Chen, M., Nurmohamed, S., Jongstra-Bilen, J., and Cybulsky, M.I. (2010). Resident intimal dendritic cells accumulate lipid and contribute to the initiation of atherosclerosis. Circ Res 106, 383–390.

Rademakers, T., Manca, M., Orban, T., Jin, H., Frissen, H., Rühle, F., Hautvast, P., Sikkink, C., Peutz-Kootstra, C., and Heeneman, S. (2018). Endothelial beta-2 spectrin: a critical plaque stiffness dependent regulator of microvessel leakage in human atherosclerotic plaque. Atherosclerosis 275, e129.

Ridone, P., Pandzic, E., Vassalli, M., Cox, C.D., Macmillan, A., Gottlieb, P.A., and Martinac, B. (2020). Disruption of membrane cholesterol organization impairs the activity of PIEZO1 channel clusters. Journal of General Physiology 152.

Saotome, K., Murthy, S.E., Kefauver, J.M., Whitwam, T., Patapoutian, A., and Ward, A.B. (2018). Structure of the mechanically activated ion channel Piezo1. Nature 554, 481–486.

Schwarz, G., Callewaert, G., Droogmans, G., and Nilius, B. (1992). Shear stress-induced calcium transients in endothelial cells from human umbilical cord veins. The Journal of physiology 458, 527–538.

Shaul, P.W., Smart, E.J., Robinson, L.J., German, Z., Yuhanna, I.S., Ying, Y., Anderson, R.G., and Michel, T. (1996). Acylation targets emdothelial nitric-oxide synthase to plasmalemmal caveolae. J Biol Chem 271, 6518–6522.

Sheetz, M.P., Schindler, M., and Koppel, D.E. (1980). Lateral mobility of integral membrane proteins is increased in spherocytic erythrocytes. Nature 285, 510–512.

Sheetz, M.P., and Singer, S. (1977). On the mechanism of ATP-induced shape changes in human erythrocyte membranes. I. The role of the spectrin complex. The Journal of cell biology 73, 638–646.

Shen, J., Luscinskas, F.W., Connolly, A., Dewey Jr, C.F., and Gimbrone Jr, M. (1992). Fluid shear stress modulates cytosolic free calcium in vascular endothelial cells. American Journal of Physiology-Cell Physiology 262, C384–C390.

Simionescu, N., Simionescu, M., and Palade, G.E. (1981). Differentiated microdomains on the luminal surface of the capillary endothelium. I. Preferential distribution of anionic sites. J Cell Biol 90, 605–613.

Sinha, B., Koster, D., Ruez, R., Gonnord, P., Bastiani, M., Abankwa, D., Stan, R.V., Butler-Browne, G., Vedie, B., Johannes, L., et al. (2011). Cells respond to mechanical stress by rapid disassembly of caveolae. Cell 144, 402–413.

Stankewich, M.C., Cianci, C.D., Stabach, P.R., Ji, L., Nath, A., and Morrow, J.S. (2011). Cell organization, growth, and neural and cardiac development require αII-spectrin. Journal of cell science 124, 3956–3966.

Tarbell, J.M., and Ebong, E.E. (2008). The endothelial glycocalyx: a mechano-sensor and -transducer. Science signaling 1, pt8–pt8.

Tzima, E., Irani-Tehrani, M., Kiosses, W.B., Dejana, E., Schultz, D.A., Engelhardt, B., Cao, G., DeLisser, H., and Schwartz, M.A. (2005). A mechanosensory complex that mediates the endothelial cell response to fluid shear stress. Nature 437, 426–431.

Uematsu, M., Ohara, Y., Navas, J.P., Nishida, K., Murphy, T., Alexander, R.W., Nerem, R.M., and Harrison, D.G. (1995). Regulation of endothelial cell nitric oxide synthase mRNA expression by shear stress. American Journal of Physiology-Cell Physiology 269, C1371–C1378.

Voas, M.G., Lyons, D.A., Naylor, S.G., Arana, N., Rasband, M.N., and Talbot, W.S. (2007). αII-spectrin is essential for assembly of the nodes of Ranvier in myelinated axons. Current biology 17, 562–568.

Wang, S., Iring, A., Strilic, B., Juárez, J.A., Kaur, H., Troidl, K., Tonack, S., Burbiel, J.C., Müller, C.E., and Fleming, I. (2015). P2Y 2 and G q/G 11 control blood pressure by mediating endothelial mechanotransduction. The Journal of clinical investigation 125, 3077–3086.

Weinbaum, S., Tarbell, J.M., and Damiano, E.R. (2007). The structure and function of the endothelial glycocalyx layer. Annu Rev Biomed Eng 9, 121–167.

Xu, K., Zhong, G., and Zhuang, X. (2013). Actin, spectrin, and associated proteins form a periodic cytoskeletal structure in axons. Science 339, 452–456.

Yamamoto, K., Korenaga, R., Kamiya, A., and Ando, J. (2000). Fluid shear stress activates Ca2+ influx into human endothelial cells via P2X4 purinoceptors. Circulation research 87, 385–391.

